# The circadian clock is required for rhythmic lipid transport in the *Drosophila* hemolymph in interaction with diet, photic condition and feeding

**DOI:** 10.1101/2023.01.24.525412

**Authors:** Kelechi M. Amatobi, Ayten Gizem Ozbek-Unal, Stefan Schäbler, Peter Deppisch, Charlotte Helfrich-Förster, Martin J Mueller, Christian Wegener, Agnes Fekete

**Affiliations:** Pharmaceutical Biology, Julius-von-Sachs-Institute, Biocenter, University of Würzburg, Julius-von-Sachs-Platz 2, 97082 Würzburg, Germany; Neurobiology and Genetics, Theodor-Boveri-Institute, Biocenter, Würzburg Insect Research (WIR), University of Würzburg, Am Hubland, 97074 Würzburg, Germany

**Keywords:** hemolymph lipids, lipidomics, circadian rhythm, feeding, locomotor activity, light-driven metabolism

## Abstract

Modern lifestyle often is at odds with endogenously driven rhythmicity, which can lead to circadian disruption and metabolic syndrome. One signature for circadian disruption is a diminished or altered cycling of metabolites in the circulating tissue reflecting the current metabolic status. *Drosophila* is a well-established model in chronobiology, but day-time dependent variations of transport metabolites in the fly circulation are poorly characterized. Here, we sampled fly hemolymph throughout the day and analysed diacylglycerols (DGs), phosphoethanolamines (PEs) and phosphocholines (PCs) using LC-MS. In wildtype flies kept on sugar-only medium under a light-dark cycle, all transport lipid species showed a synchronized bimodal oscillation pattern with maxima at the beginning and end of the light phase which were impaired in *period^01^* clock mutants. In wildtype flies under constant dark conditions, the oscillation became monophasic with a maximum in the middle of the subjective day. In strong support of clock-driven oscillations, levels of DGs, PEs and PCs peaked once in the middle of the light phase under time-restricted feeding independent of the time of food intake. Rearing of wildtype flies on lipid-containing standard medium masked the rhythmic alterations of hemolymph lipid levels. Our data suggest that the circadian clock aligns daily oscillations of DGs, PEs and PCs in the hemolymph to the anabolic siesta phase, whith a strong influence of light on phase and modality. This finding opens the question whether and to what extent the circadian regulation of transport lipid levels in the hemolymph contributes to the health of the fly.

## Introduction

Physiological processes need to be temporally aligned with each other and with the environment to ensure health and well-being of organisms. In animals, including humans, circadian clocks in the brain and peripheral organs are endogenous timers that are vital for temporal alignment across the body even in the absence of environmental synchronisation cues (Zeitgeber such as light or feeding) (1–3). Moreover, circadian clocks allow animals to prepare their physiology and behaviour in anticipation of periodic biotic and abiotic changes in the surrounding world (4). Circadian disruption and misalignment to daily environmental changes promote diverse physiological pathologies, including metabolic disorders such as type-2 diabetes and obesity in shift workers or people with sleep disorders (5, 6).

Like other metabolic pathways, lipid metabolism shows daily rhythmicity and is under circadian regulation in mammals (see (7, 8)). In particular, there is substantial daily rhythmicity in transport lipids in the plasma, including di- and triacylglycerols (DGs, TGs), free fatty acids, sphingolipids and phospholipids (e.g. (9–12)). The fruit fly *Drosophila*, a long-standing model in circadian biology, is increasingly attracting attention as a model to study lipid metabolism in health and disease (13), including circadian aspects (14). Global metabolic profiling in *Drosophila* identified a larger set of molecules including acyl carnitines, fatty acids and other lipids to rhythmically oscillate in a light- and clock-dependent manner (15–17). Moreover, transcripts for a large set of lipid-metabolising enzymes are cycling in a circadian fashion in the gut and fat body (14, 18), main organs in the regulation of systemic lipid metabolism in the fly. These enzymes are responsible for different chemical steps in lipid metabolism, including long-chain fatty acid metabolism, fatty-acyl-CoA reduction, TG/DG breakdown and synthesis as well as β-oxidation.

Unlike in mammals, diel temporal patterns of transported lipids in the *Drosophila* hemolymph (the fly “blood”) remain largely uncharacterised, and their dependency on the circadian clock is unclear. To fill this gap, we profiled the diel and circadian temporal pattern of lipids in the hemolymph of *Drosophila* using ultra performance liquid chromatography coupled to time-of-flight mass spectrometry (UPLC-TOF-MS) in Canton-S wildtype (WT_CS_) flies and *period^01^* mutants (*per^01^*) with a defective molecular clockwork. We focused on the two major groups of transported lipids: glycerolipids (DG) and phospholipids (phosphoethanolamines (PEs), phosphocholines (PCs)). DGs were chosen as they represent the transport form of fatty acids used for cellular energy and lipid homeostasis in insects (19, 20), and since TG/DG breakdown and synthesis is under transcriptional control of the fat body clock (18). Phospholipids play a major role as components of biological membranes, in signal transduction and in membrane trafficking, and serve as building blocks for more complex cellular lipids. DGs, PEs and PCs are main lipid constituents of Lipophorins, the major hemolymph carrier vehicle for water-insoluble metabolites (20, 21).

As the temporal profile of lipid metabolism is strongly influenced by time-restricted feeding (TRF) in flies and mammals (e.g. (22–25)), we performed hemolymph lipid profiling in flies kept on different diets or TRF regimes. Our findings show that the circadian clock, light and feeding are major factors that shape the temporal profile of circulating lipids in flies. Our results further provide evidence for a role of *de-novo* lipid synthesis in clock-dependent temporal changes of transport lipids in the hemolymph. Interestingly, the clock seems to align peak levels of transport lipids to the anabolic siesta time, which may have a systemic and health-relevant impact.

## Materials and methods

### Fly husbandry

WT_CS_ and *per^01^* flies were a kind gift of Bambos Kyriacou (Leicester). Flies were raised on standard *Drosophila* medium consisting of 0.8% agar, 2.2% sugar beet syrup, 8.0% malt extract, 1.8% yeast, 1.0% soy flour, 8.0% corn flour and 0.3% hydroxybenzoic acid under a 12:12 h light:dark regime (LD) at 25±0.2 °C and 60±2% relative humidity. For the experiments, young males were collected within 24 hr after eclosion and transferred to vials containing either standard medium or sugar-only medium consisting of 5% sucrose and 2% agar.

### Hemolymph sampling

Six days old male flies were transferred from their food vials into a clean vial and anesthetized on ice for 3 min. Then the males were placed on a plastic petri-dish wrapped with parafilm to reduce water condensation under a stereomicroscope. A small incision was made on the thorax of the flies using a sharp tungsten needle. After incision, 20 male flies were pooled into a 0.5 ml Eppendorf tube with three small holes at the bottom, which was inserted into a 1.5 ml Eppendorf tube (Suppl Fig S1). Subsequently, tubes with the flies were centrifuged for 5 min at 4 °C at 3075 rcf (5000 rpm) in a benchtop centrifuge. After centrifugation, clear and light-yellowish hemolymph samples were typically obtained. Cloudy deep-yellow samples were discarded. The obtained hemolymph in the 1.5 ml Eppendorf tube was then taken up with 0.5 or 1 μl micro capillaries. The length occupied by the hemolymph in the micro-capillaries was measured under a stereomicroscope and the volume was calculated. Afterwards, the microcapillaries were each placed in an Eppendorf tube containing 25 μl of Millipore water and all its content ejected using a small rubber ejector. The test tubes containing samples were stored at −80 °C until sample preparation for lipid measurement.

### Sample preparation and lipid analysis

Prior to analysis, the diluted hemolymph samples were dried in a rotating evaporator at 50 °C until complete dryness and reconstituted in 75 μl of cold isopropanol containing internal standards (1 ng/μl of DG(20:0), PE(34:0) and PC(34:0) and 0.1 ng/μl of TG(30:0)), followed by ultrasonication for 15 minutes. Analyses were performed using an Acquity Ultra Performance LC coupled to a Synapt G2 HDMS (Waters) as previously described (16). Briefly, a BEH C18 column (2.1×100 mm, 1.7 μm, Waters) at 60 °C was used for chromatographic separation. A linear binary solvent gradient was applied using 30-100 % eluent B over 10 min at a flow rate of 0.3 ml/min. Eluent A consisted of 10 mM ammonium acetate in 60/40% water/acetonitrile and eluent B consisted of 10 mM ammonium acetate in 90/10% isopropanol/acetonitrile. After chromatographic separation, metabolites were ionized using an electrospray ionization (ESI) source operated in positive mode. The quadrupole was operated in a wide-band mode, and data was acquired over the mass range of 50-1200 Da in a centroid data mode. Leucine-enkephaline (m/z 556.2771) was used for internal calibration every 0.3 min. MassLynx and QuanLynx (version 4.1; Waters) were used to acquire and process chromatograms. For quantitation, peak areas of the analytes and internal standards (IS) in the extracted ion chromatogram (XIC) were integrated. Concentrations were calculated using an internal standard technique (response factor of one) for each analyte/IS pair and plots were generated using either R or MS-Excel. The retention time and mass-to-charge ratios of the lipids are listed in Suppl Table S1.

### Locomotor activity recording and feeding assay

*Drosophila* Activity Monitors (DAM-2, Trikinetics, Waltham, USA) were used for recording locomotor activity. For both feeding conditions (standard medium or sugar-only medium), 32 male flies per genotype (WT_CS_ and *per^01^*) were individually placed in small glass tubes (5 mm diameter) which contained food on one side closed with a rubber plug. The other side was closed with a small foam plug to allow for air exchange. Standard medium consisted of 3.6% yeast extract, 5% sucrose and 2% agar. Sugar-only medium consisted of 2% agar containing 4% sucrose. Flies were monitored under LD12:12 (LD) conditions for seven days, and constant darkness (DD) for the rest of the experiments at constant humidity (60%) and temperature (25 °C). Activity of the flies was measured by counting the interruptions of the infrared beam in the middle of the tubes.

Capillary feeding (CAFE) assay was performed as previously described (16). Standard medium contained 3.6% yeast extract and 5% sucrose, while sugar-only medium consisted of 5% sucrose. Both media contained 0.3% FD&C Blue No 1 (E133) as coloring agent.

### Statistical analysis

Time-dependent lipid oscillations and food consumption were analyzed by JTK_CYCLE in an R environment (26). To identify oscillation profiles, period length was set to 6-24 h. Lipids were regarded to be oscillating if the adjusted p-value (adj.p) was less than 0.05. JTK_CYCLE also calculated amplitude (AMP) and time of maximum aka lag phase (LAG). Rhythmicity of locomotor activity was analysed using Lomb–Scargle and Chi-squared test in ActogramJ (27).

## Results

### Diacylglycerol and phospholipid levels change over the day in a clock- and light-dependent fashion on sugar-only medium

Previous studies have reported daily oscillations of several lipids in fly bodies (thorax and abdomen) or in whole flies (15–17), but it remained unclear whether lipid levels in the fly hemolymph vary according to the time of day. To fill the gap, we analysed lipids in extracted hemolymph every 3h over the entire day in six days old male WT_CS_ flies kept under a LD12:12 (LD) photoperiod (Suppl Fig S2a) and fed *ad-libitum* with sugar-only medium which contained sugar as the sole carbon- and energy source. First, we profiled the major hemolymph lipid species (DGs, PEs and PCs) using UPLC-TOF-MS and identified 20 lipid-species (DG(26:0), DG(26:1), DG(28:0), DG(28:1), DG(30:1), DG(32:1), DG(32:2); PE(30:1), PE(32:1), PE(32:2), PE(34:1), PE(34:2), PE(34:3), PE(36:2), PE(36:3); PC(32:1), PC(32:2), PC(34:1), PC(34:2), PC(34:3) listed in Suppl Table S1). Interestingly, all lipid-species of the three different lipid classes showed very similar temporal oscillation patterns in phase with each other (Suppl Fig S3), indicating that the daily variation is independent of the lipid-species. This allowed us to sum up the total levels for each lipid class. The experiments were independently repeated seven times over a period of three years. In six out of the seven experiments, levels of total DG, PE and PC species peaked at the beginning (ZT1) and end (ZT10-13) of the light phase with variable amplitudes (Suppl Fig S4). Next to manual inspection, daily oscillations were analyzed using the non-parametric algorithm JTK_CYCLE (26) which revealed significant daily oscillations in five out of the seven individual experiments (adj.p < 0.01, Suppl Table S2). Next, we combined and averaged the normalized individual experiments which revealed a strong bimodal daily oscillations in hemolymph DGs, PEs and PCs with peaks at ZT1 and 10 (Fig 1a), which is reflected by low adj.p values (4e^-7^- 3e^-6^) shown in Suppl Table S3.

**Figure 1:**
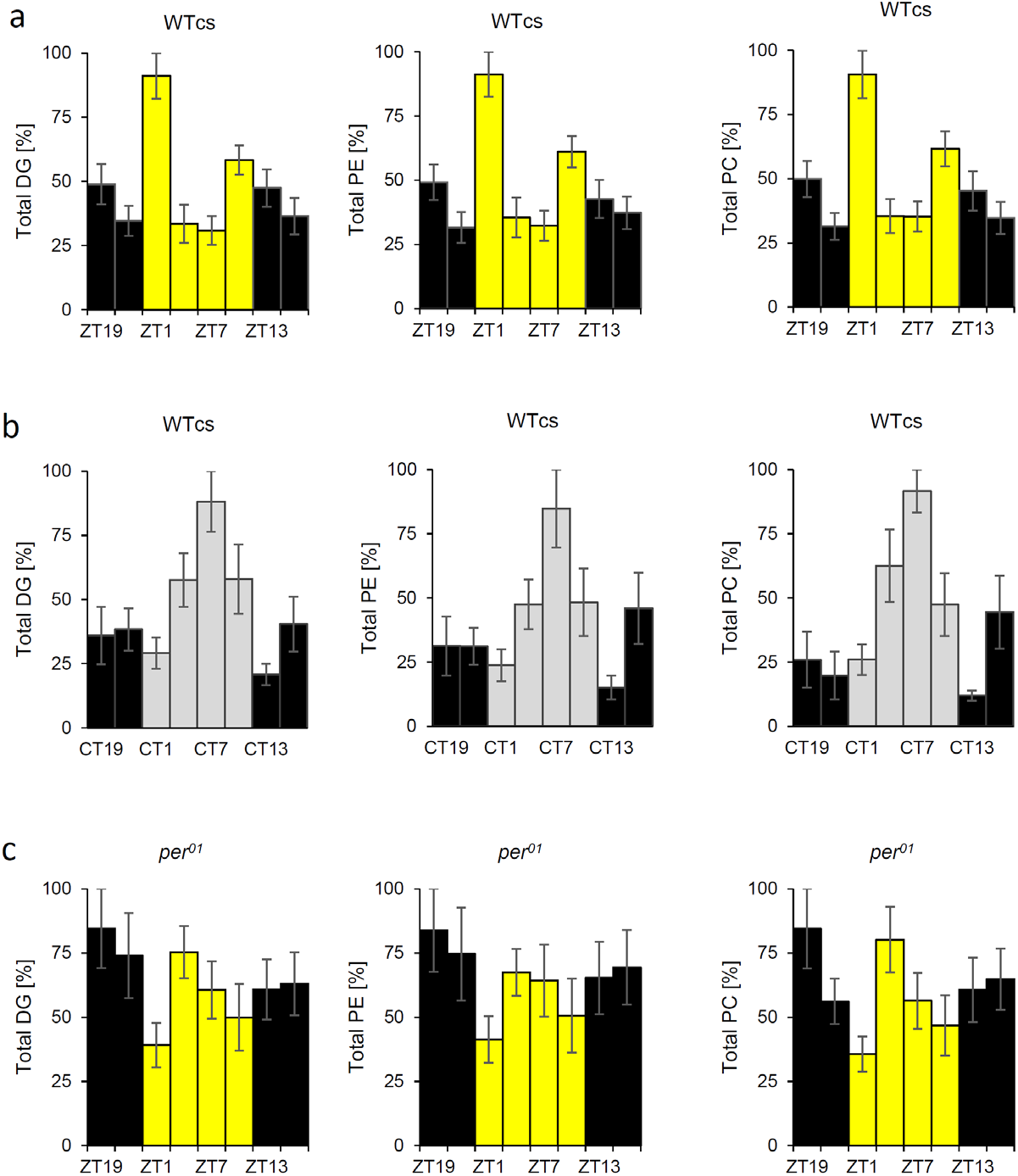
Clock and light dependent diel oscillation of hemolymph lipids. Normalized levels of DGs (left), PEs (middle) and PCs (right) in male flies kept on sugar-only medium. WT_CS_ flies in LD (a, N=7, n=91) and DD (b, N=3, n=39), and per^01^ flies in LD12:12 (c, N=3, n=39). Colour of the bars indicates light conditions. (LD: yellow = lights on, black = lights off; DD: light gray = subjective light phase, dark gray = subjective dark phase). Data represent mean ± standard error.

To test whether the observed bimodal oscillation of hemolymph lipids is endogenously driven by the circadian clock, we next analysed DGs, PEs and PCs in the hemolymph of WT_CS_ in DD. Flies were synchronized for three days in LD and switched to DD conditions before hemolymph was sampled every three hours on the sixth day under red light to which the circadian clock of the fly is blind (Suppl Fig S2b). In two out of three independently performed experiments, total levels of DGs, PEs and PCs showed a monomodal peak at subjective midday (Suppl Fig S5). However, JTK_CYCLE identified significant oscillation only in one experiment (Suppl Table S2). The combined normalized and averaged lipid levels of all three experiments showed a monomodal peak at CT7 (Fig 1b). Though profiles of DGs, PEs and PCs are very similar, JTK-Cycle revealed rhythmicity only for PCs (adj.p: 0.003 for PC, 0.2 for PE and 1 for DG, Suppl Table S3).

The difference in the temporal profile between LD and DD suggests that light has a significant impact on the daily fluctuations of hemolymph DGs, PEs and PCs in *Drosophila*. In LD, lipid levels peak bimodally around the times of lights on and off, while in DD only one peak occurs in the middle of the subjective day (CT7). Nonetheless, the monomodal oscillation in DD provides strong evidence for a circadian base underlying the daily changes of hemolymph lipids. Moreover, light seems to positively affect total lipid levels in the hemolymph of WT_CS_. Quantified lipid levels after averaging of all sampling timepoints were considerably higher in LD than in DD in two out of three experiments (fold changes of averaged levels between DD and LD: 0.2, 0.3 and 1, Suppl Table S4).

To better disentangle the effect of light and the circadian clock on the daily fluctuations of circulating lipids in *Drosophila*, we determined the levels of DGs, PEs and PCs in the hemolymph of *per^01^* clock mutants under LD (Suppl Fig S2a). As light has a significant impact on hemolymph lipid oscillations in wildtype flies, we expected a weak bimodal activity pattern in the clock-impaired flies. Yet, in contrast, daytime dependent levels of DGs, PEs and PCs in the hemolymph of *per^01^* mutants under LD varied strongly between the three independently performed experiments (Suppl Fig S6) and oscillations were only significantly rhythmic in one out of three experiments according to JTK_CYCLE (Suppl Table S2). After averaging the normalized data, levels of hemolymph lipids did not show systematic daily variations (Fig 1c) and rhythmicity was not detected (JTK_CYCLE adj.p: 0.6 and 1, Suppl Table S3). Interestingly, averaged total lipid levels in the clock null mutant were wild-type like under LD condition, with fold changes between *per^01^* and WT_CS_ of 1.0-1.4 for DG, 1.1-2.1 for PEs and 1.3-2.1 for PC (Suppl Table 4). These results suggest that a disrupted clock in *per^01^* mutants mostly affects the temporal profile of hemolymph lipid levels but not total levels. Importantly, the observed arrhythmicity of *per^01^* further shows that light by itself is not sufficient to drive rhythmicity in hemolymph lipid levels, but rather modulates the phase and shape of the clock-dependent rhythm.

### Dietary lipids mask diel lipid oscillation patterns in the fly hemolymph

The result above show that hemolymph lipids oscillate in WT_CS_ flies when diet is restricted to sugar. We next asked whether daily rhythmicity in circulating lipids in *Drosophila* persists on a lipid-containing diet. We determined levels of DGs, PEs and PCs in the hemolymph of WT_CS_ male flies fed *ad libitum* with standard medium in three-hour intervals within one day in LD (Suppl Fig S2a). On standard medium, DGs in the hemolymph can originate from the gut where they can be reassembled from dietary fatty acids or *de novo* produced from sugars, or from TG mobilization from the fat body (28). We found that the absolute levels in the hemolymph of flies kept on standard medium were about five times higher for DGs, seven times higher for PEs and doubled for PCs compared to sugar-only medium (Suppl Table S4), which is consistent with a previous study (21). The relative amounts of hemolymph DGs was biased towards medium chain lengths (26 to 28) compared to a bias towards longer chain lengths (30-34) in whole bodies or heads comprising the fat body (Suppl Fig S7). Importantly, the DG composition differed considerably between the hemolymph, head and body and the standard medium (Suppl Fig S7, head and body data from an earlier study (16)).

The temporal patterns of the individual DG, PC and PE species on standard medium were largely similar (Suppl Fig S8). This allowed us again to calculate their total levels which fluctuated only weakly with maxima between ZT22-ZT1 and at ZT13 in three out of six independently performed experiments (Suppl Fig S9, Suppl Table S2). This weak bimodal peak pattern (Fig 2) showed a relative increase of the evening peak compared to the situation on sugar-only medium (Fig 1a). However, after normalisation and averaging of all six experiments, JTK_CYCLE did not detect a significant daily rhythmicity (adj.p values = 0.6 and 1.0 shown in Suppl Table S3). Arguably, the most likely reason for the loss of hemolymph lipid oscillations on standard medium is masking by the consumption of exogenous dietary lipids. Collectively, the results suggest that daily rhythmicity in the hemolymph profiles of the analysed lipids is mainly determined by mobilization from intracellular stores in the fat body or *de novo* synthesis from the gut, while flies stabilize hemolymph transport lipids at a constant high level if they have *ad libitum* access to dietary lipids. These levels may mask or impair the effect of rhythmic lipid *de novo* synthesis or mobilisation from intracellular stores.

**Figure 2:**
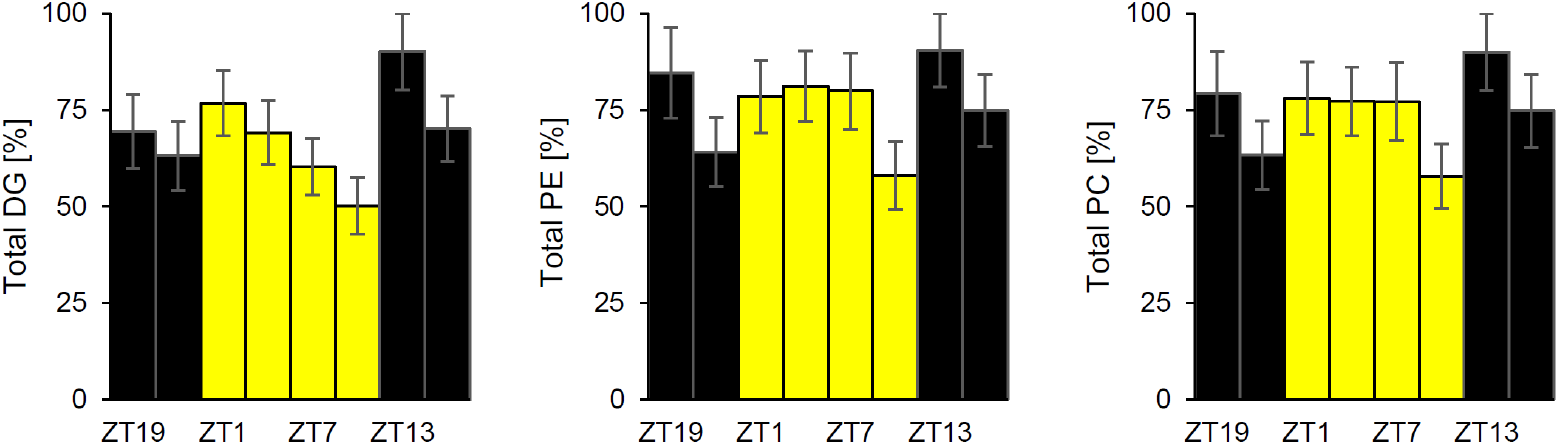
Dietary lipids mask diel rhythmicity in hemolymph lipids. Normalized total levels of DGs (left panel), PEs (middle panel) and PCs (right panel) in the hemolymph of WT_CS_ male flies on standard medium. Colour of the bars indicates light conditions (yellow = lights on and black = lights off). Data represent mean ± standard error (N=6, n=78).

### Feeding and locomotor activity rhythmicity are not phase-coupled to diel lipid oscillations under lab conditions

Feeding and locomotor activity are closely associated with metabolism and are modulated by the circadian clock in *Drosophila* (29–31). Therefore, we asked to which extent both behaviors contribute to the temporal profile of hemolymph lipids. We employed a CAFE assay (32) to measure feeding, and TriKinetics activity monitors to record locomotor activity of WT_CS_ and *per*^01^ male flies. In line with our previous work (16), we observed a significant diel rhythmicity in locomotor activity (Suppl Fig S10a) and food consumption (Suppl Fig S11a) in WT_CS_ on standard medium under LD (Fig 3a). In DD, circadian rhythmicity of locomotor activity (Suppl Fig S10a) and food consumption (Suppl Fig S11a) persisted, but with lower amplitude (adj.p food consumption: 6e^-18^ in LD, 1e^-6^ in DD; % flies with rhythmic locomotor activity: 97% in LD, 97% in DD, Suppl Table 5). In contrast, the rhythmicity of *per^01^* mutants was already low in LD (adj.p food consumption: 4e^-3^; 81% flies with rhythmic locomotor activity, Suppl Table 5) and became arrhythmic in DD (Suppl Fig S10b, S11b).

**Figure 3:**
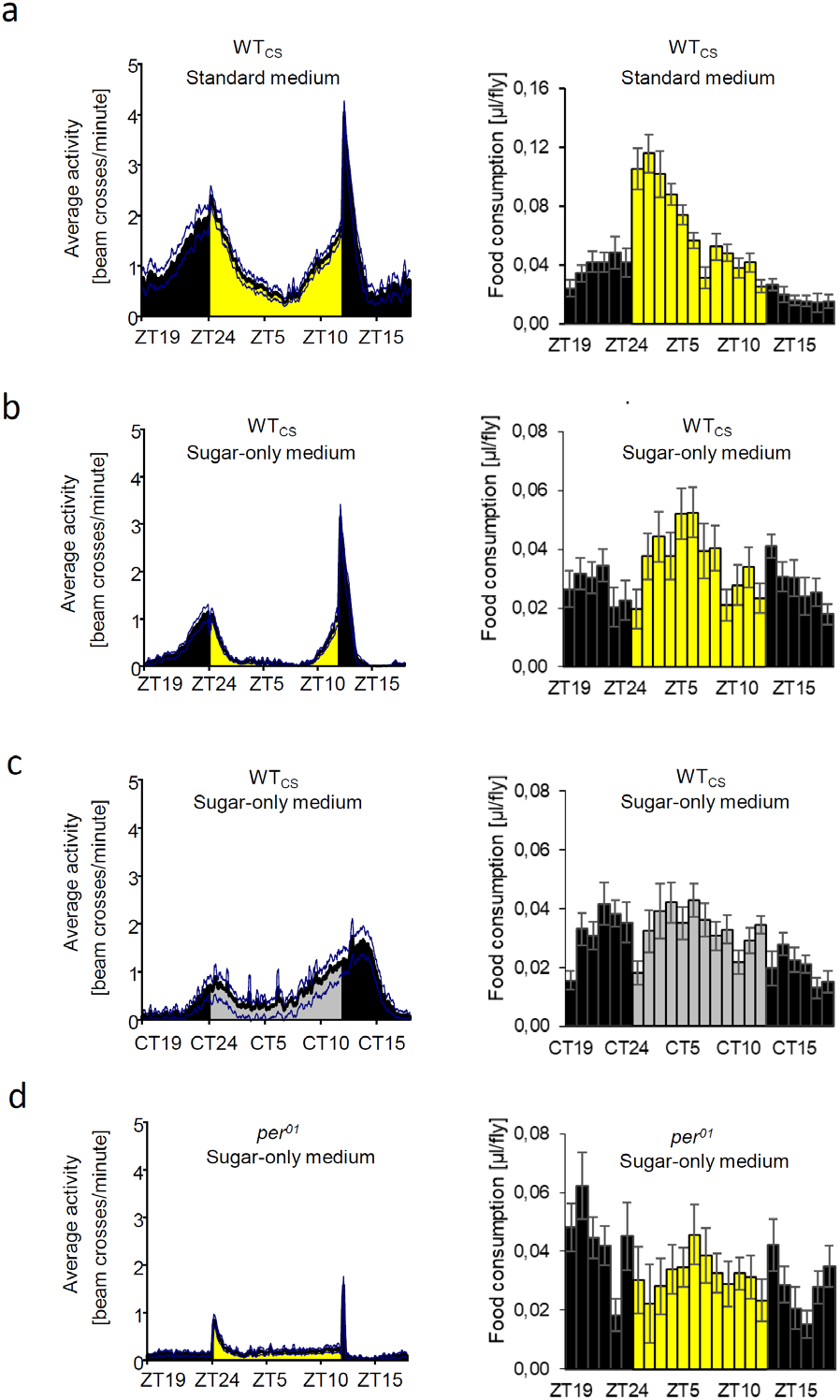
Diel rhythmicity profiles of locomotor activity and food consumption differ in WT_CS_ flies. Locomotor activity (left) and food consumption (right) were monitored in WT_CS_ in LD on standard medium (a) and sugar-only medium (b). In addition, both behaviours were investigated in WT_CS_ in DD (c) and per^01^ in LD (d) on sugar-only medium. Colour of the bars indicates light conditions (LD: yellow = lights on and black = lights off; DD: light gray = subjective light phase; dark gray = subjective dark phase). Data represent mean ± standard error (N=2, n=10 for CAFÉ assays and n=29-32 for locomotor activity).

Next, we asked whether a sugar-only diet alters the rhythmicity in feeding and locomotor activity in WT_CS_ flies. On sugar-only medium, the bimodal pattern of locomotor activity (Suppl Fig S12a) persisted under LD (Fig 3b) and DD conditions (Fig 3c) in WT_CS_ (91% rhythmic flies in LD and 78% rhythmic flies in DD, Suppl Table 5). This is not surprising as many *Drosophila* chronobiology labs routinely use sugar-only medium during locomotor activity recording. However, diel oscillation in food consumption in WT_CS_ strongly declined on sugar-only medium in LD (Fig 3a,b). JTK_CYCLE analysis of the normalized data of food consumption on each diet confirmed the reduced feeding rhythmicity on sugar-only medium compared to standard medium (adj.p in sugar-only medium: 1e^-3^, standard medium: 1e^-18^, Suppl Table 5), and revealed a phase shift by around 4 hr (maximum (LAG) in sugar-only medium: ZT5, standard medium: ZT1, Suppl Table 5). Manual inspection of food consumption revealed dampened oscillations in WT_CS_ in DD (Fig 3c) compared to LD (Fig 3b), with similar oscillation parameters of the normalised data (adj.p: 1e^-3^ in LD, 2e^-5^ in DD; LAG in LD and DD: ZT5 and CT5; relative amplitude: 13% in LD, 11% in DD, Suppl Table S5). The rhythmicity in food consumption on sugar-only medium seems to be under control of the circadian clock as feeding behaviour in *per^01^* clock mutants was arrhythmic in LD (Fig 3d, adj.p: 0.6-1, Suppl Table 5) as well as in DD (Suppl Fig S13b).

On standard medium, the peaks of food consumption and locomotor activity of WT_CS_ flies (Fig 3a) did not correlate with peaks in hemolymph lipids (Fig 2). On sugar-only medium in LD, however, the peaks of both behaviors (Fig 3b) temporally coincided with peaks in the circulating lipid titer (Fig 1a), suggesting that light synchronizes locomotor activity, feeding and circulating lipid levels in absence of dietary lipids. This synchronisation was lost in DD, as the phases of locomotor activity, food consumption (peaks around dusk and dawn, Fig 3c) and hemolymph lipids (peak at midday siesta, Fig 1b) separated. Specifically, locomotor activity peaked around CT0 and CT12 whereas the maximum of feeding behaviour was CT5 (Suppl Table 5) and hemolymph lipid titers peaked between CT7 and CT9 (Suppl Table 3). This suggests that, under our conditions, the observed peaks in hemolymph lipids are not a consequence of activity-induced lipid mobilisation.

### The time of restricted feeding does not alter the phase of hemolymph lipid oscillations

To more directly test whether lipid oscillations are a consequence of food consumption, we used a time-restricted feeding (TRF) paradigm. Flies were fed sugar-only medium only during the 12 hr light phase or the 12 hr dark phase. First, we monitored food consumption in WT_CS_. During light phase-TRF, flies showed a tendency to feed more between ZT3 and ZT6 (Fig 4a) but consumption was more or less stable throughout the light phase. During dark phase-TRF, food consumption was more variable without a consistent pattern during the 12 hr of darkness (Fig 5a). Switching the flies back from TRF to *ad libitum* feeding resulted in a normal diel rhythmicity of food consumption on the second day (Fig 4b, Suppl Table 5) as previously reported (33).

**Figure 4:**
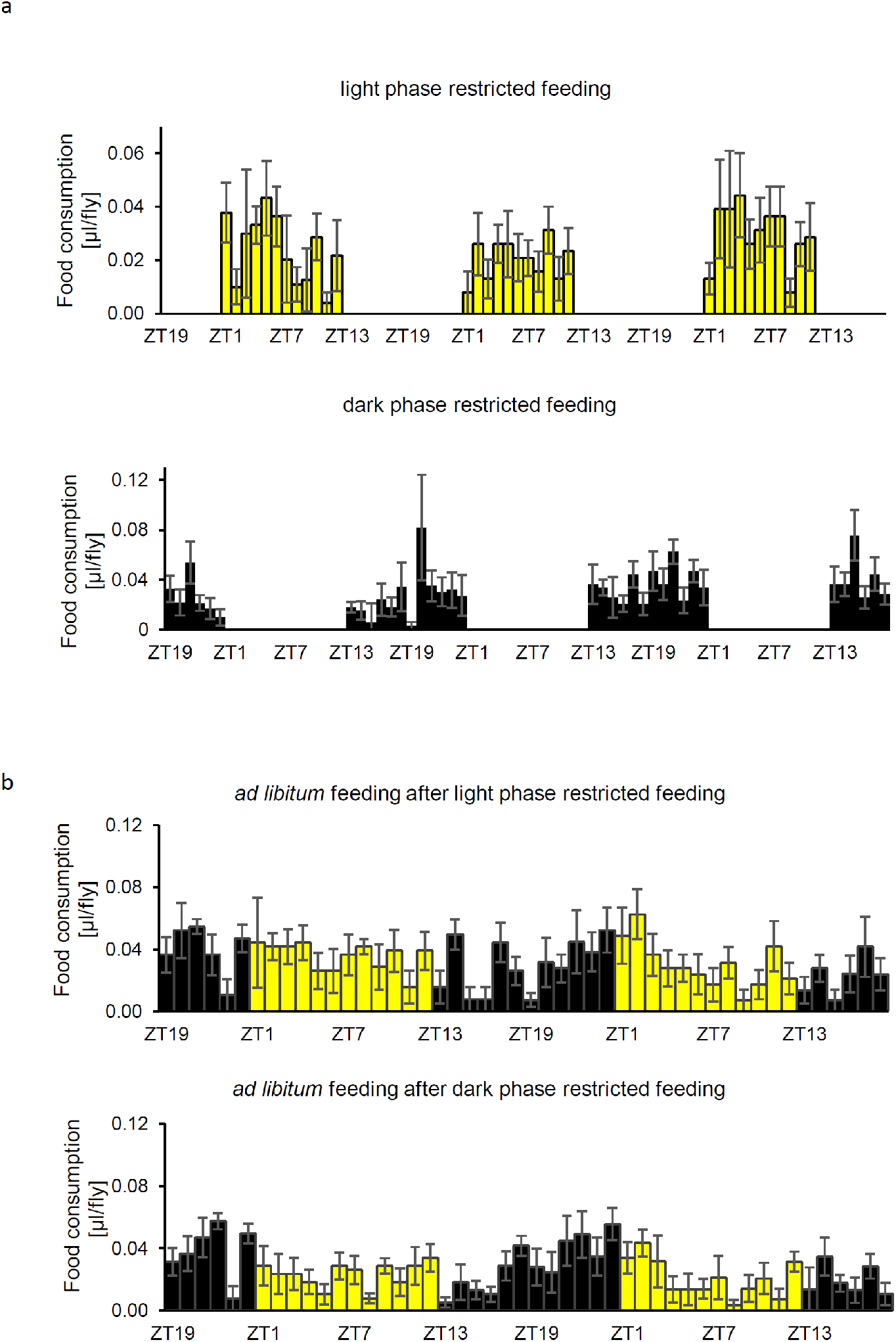
Time-restricted feeding does not entrain the temporal pattern of food consumption. WT_CS_ males were fed only during the light ((a) top) or dark phase ((a) bottom) in LD. Food consumption was monitored from the fourth to sixth day using a modified CAFÉ assay. After TRF, flies were fed ad libitum with sugar-only medium and food consumption in LD was monitored for two further days (b). Colour of the bars indicates light conditions (yellow = lights on and black = lights off). Data are presented as mean ± standard error (n = 5).

**Figure 5:**
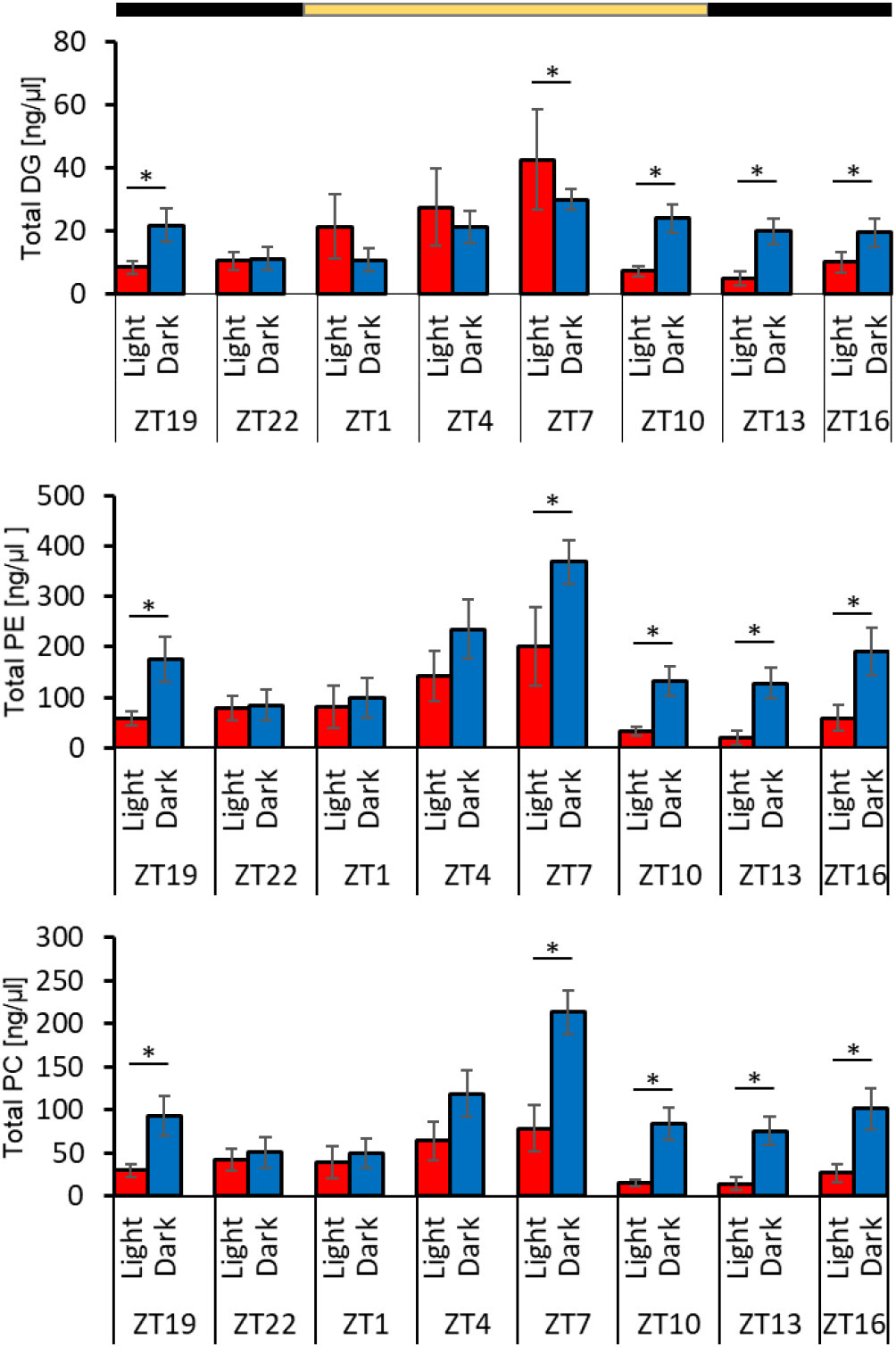
Time-restricted feeding does not alter the phase of hemolymph lipid oscillations. WT_CS_ males were fed for three days only during the light (red bars) or dark phase (blue bars) in LD, and total levels of DGs (top panel), PEs (middle panel) and PCs (bottom panel) in the hemolymph were measured. Asterisks denote statistically significant differences between light phase-TRF and dark phase-TRF (p < 0.05, t-test). The bar at the top indicates lights-on (yellow) and lights-off (black). Data are presented as mean ± standard error (n= 11).

Next, we analyzed the levels of DGs, PEs and PCs in LD in the hemolymph of WT_CS_ flies subjected to light phase- or dark phase-TRF for three consecutive days. Hemolymph was sampled every three hours on the last day of TRF (Suppl Fig 2c). Independent of the TRF conditions, levels of hemolymph lipids oscillated unimodally with a maximum at ZT6-ZT10 (Fig 5). Main differences in the pattern pertained to the amplitude at ZT7, which was significantly higher for DGs and significantly lower for PEs and PCs in light phase-TRF than in dark phase-TRF. Moreover, the levels of all three lipid classes were constantly higher from ZT10-ZT19 in dark phase-TRF flies compared to light phase-TRF flies. These differences correlate well with the time when flies had access to food. Time-dependent levels of DGs, PEs and PCs in the hemolymph were not always classified as rhythmic by JTK_CYCLE possibly due to the high variance leading to high adj.p value (Suppl Table 3). Interestingly, oscillation profiles of WT_CS_ hemolymph lipids under TRF in LD were very similar to the profile observed in *ad libitum* feeding in DD, with a monomodal peak around midday. In addition, the finding of a monomodal peak in lipid levels around midday independent on whether TRF was limited to the photo- or scotophase (Fig 5) provides strong evidence that the temporal profile of hemolymph lipid levels is under control of the circadian clock and is not a direct consequence of feeding.

## Discussion

The fruit fly *Drosophila* is a prime model in circadian research (34), and offers ample opportunity to study the interplay between the circadian clock and metabolism. Consequently, several metabolomic studies have characterised the global effects of an impaired clock on metabolism in *Drosophila*, including lipids (15–17). Nevertheless, the temporal profile and clock dependency of transport lipids in the hemolymph remained unknown. Transport lipids represent a hub for lipid mobilisation and *de novo* synthesis and can serve as a proxy to assess overall lipid metabolism. Therefore, we here characterised the diel profile of the major lipid classes in the hemolymph of the fruit fly *Drosophila* and its dependency on the circadian clock, light, locomotor activity and feeding.

### Lipids in the fly hemolymph show diel oscillations that are influenced by the clock and light and masked by dietary lipid intake

A major conclusion from our work is that hemolymph DG, PE and PC levels show clock- and light-dependent oscillations in phase with each other in flies kept on sugar-only diet. Specifically, levels of DGs, PEs and PCs peaked in the early morning and late afternoon in the hemolymph of male WT_CS_ in LD on sugar-only medium. This temporal profile was altered in DD to a monomodal oscillation with peak in the subjective midday, while oscillations were strongly impaired in LD in *per^01^* clock mutants. Taken together, the results suggest that the circadian clock is the main driver behind the oscillations, while light exerts a significant effect on the phase and temporal profile. These findings are in line with metabolomic studies on heads and bodies of *Drosophila*, which also found an influence of both circadian clock and the light phase on the oscillations of lipids and other metabolites (15, 16).

Noteworthy, *per^01^* clock mutant flies showed unaltered total levels of circulating lipids in LD in our study. This is in contrast to the situation in rodents where disruption of the molecular clock lead to altered levels of circulating lipids (see (35)). For example, levels of plasma TGs were higher in *Bmal1* (36) and lower in *Per2* mutant mice (37).

Another interesting finding of our work is that DGs, PEs and PCs levels in the hemolymph oscillate in phase with each other. This result is at odds with the situation in humans in which the daily oscillation profiles of glycerolipids and PCs in the blood are not in sync on a lipid-poor diet (9). This difference may be easily explained by differences in the mode of lipid transport between insect and mammals. In mammals, the inter-organ transport of lipids occurs by multiple and organ/lipid species-specific means, including chylomicrons and lipoproteins, and the timing of circulating TGs is dependent on the type of lipoprotein (high density lipoproteins or apoB-lipoprotein) (38). In the insect hemolymph, DGs are primarily transported via lipophorin particles that are surrounded by phospholipids including PEs and PCs that originate from the fat body (20, 21). We assume that this co-shuttling of glycero- and phospholipids between organs in lipophorin particles underlies the observed phase synchronicity of circulating lipids in flies.

Diet impacts the circadian clock machinery (39) and diet was shown to affect metabolite oscillations as well (40–42). In mice, for example, the majority of serum lipids loose diel rhythmicity after a high fat diet (40). Our study finds that hemolymph lipid oscillations were strongly dampened and disappear when flies had *ad libitum* access to a rich standard food containing lipids, sugars and proteins as well as micronutrients. We conclude from this that continuous access to exogenous (dietary) supply of lipids and other macronutrients attenuates and masks the diel oscillations of glycero- and phospholipids in the hemolymph of *Drosophila*, most likely via dietary intake. The overall higher levels of hemolymph lipids in flies on a lipid-containing diet compared to sugar-only medium is in line with earlier data (21, 43) and might be attributed to increased intake of dietary fat. Collectively, our data points towards an overall dominant influence of dietary lipid intake on lipid levels in the fly hemolymph.

The composition of DGs in our standard food (biased toward long chain fatty acids) was clearly different from the glycerolipid composition in the hemolymph (biased towards medium chain fatty acids). This suggests that dietary DGs are not directly transported into the hemolymph, but are metabolised or modified by the insect digestive tract. This observation is in agreement with earlier data that suggest a difference in DG species due to shortening of dietary fatty acids prior to export to target organs via the hemolymph (43).

### Rhythmicity of DGs in the fly hemolymph on sugar-only medium likely involves de-novo synthesis in the gut

The primary sources of hemolymph lipids on sugar-only medium are either mobilisation from the fat body, or *de-novo* synthesis from the gut (see (19, 21, 44)). We did not investigate this further here, though it could be done by stable isotope-labeled feeding. Nevertheless, two findings suggest that lipid *de-novo* synthesis is daytime-dependent at least on a nutrition-depleted diet. First, the peak times of hemolymph lipid levels and locomotor activity (the dominant energy-consuming behaviour possible under our laboratory conditions) are not synchronized or in a fixed phase-relationship (as would be expected if mobilisation is involved), but dissociate in DD. Second, we observed a bias towards medium-chain DGs (DG26:X, DG28:X) and a smaller degree of desaturation for DGs in the hemolymph than in whole heads or bodies comprising the fat body. Although, in absolute levels, the majority of DGs is found in the hemolymph, they can also be found in the fat body (43). In the hemolymph, medium-chain DGs are by far the dominant DG species while long chain DGs constitute only a very small fraction. In contrast, the fat body contains about similar levels of DGs with medium and long-chain fatty acyls (43). The medium-chained DGs are mostly produced in the gut, as their levels in the gut are massively increased upon genetic downregulation of lipophorin (*Lpp*) (21). This effect is also seen on lipid-depleted medium (21).

### Diel oscillations in hemolymph lipids are not driven by rhythmic locomotor activity or feeding

Metabolism is strongly influenced by physical activity and nutrient uptake. To dissect their influence on the diel hemolymph lipid oscillations we compared the temporal lipid profiles with that of the energy supply behaviour (feeding) and the most overt energy-consuming behaviour (locomotor activity). On standard medium in LD, hemolymph lipid levels maintained a relatively constant level while both feeding behaviour and locomotor activity oscillated with distinct morning and smaller evening peaks. This suggests that energy supply and expenditure are at equilibrium under these conditions. In addition, both feeding behaviour and locomotor activity obviously do not exert a direct instantaneous influence on hemolymph lipid oscillations. This seems also to hold true on sugar-only medium, where lipid levels were out of phase with feeding (LD) or locomotor activity (DD). Further support comes from the finding that *per*^01^ mutants in LD lost their rhythmic diel oscillations in hemolymph lipids and feeding behaviour although locomotor activity maintained light-driven rhythmicity.

We note that our experiments were carried out with flies kept in small vials offering little possibility and incentive for intense and sustained physical exercise that requires lipid mobilisation. It would thus be interesting to repeat similar experiments under a more natural setting which permits energy-craving activities such as intense foraging or flying.

In mammals and *Drosophila* alike, TRF is a strong “Zeitgeber” that is able to reset the circadian clock (23, 45–49). Changes in feeding pattern or feeding time with different TRF feeding paradigms influences hepatic lipid homeostasis in mice (50, 51). Also, in lactating Holstein cows, 8-12 hours shift in peaks of metabolite levels, plasma hormones and milk synthesis were observed between nighttime and daytime restricted feeding (52). Here, we found that both light phase-TRF and dark phase-TRF shifted the hemolymph lipid peak to the middle of the light phase when compared to *ad libitum* feeding on sugar-only medium. TRF thus does not entrain but phase-shift the rhythmicity of circulating glycero(phospho)lipids in *Drosophila*, an effect strikingly similar to the situation for hepatic TGs in mice (50).

In conclusion, our data show that DGs, PEs and PCs in the fly hemolymph oscillate in a clock- and light-dependent manner in flies without access to dietary lipids. This cycling appears largely independent of the time of feeding. Based on the observed differences in lipid species composition between the hemolymph, fat body and food source, we suggest that the observed lipid oscillations on lipid-free diet is primarily due to rhythmic *de novo* lipid synthesis in the gut, and not rhythmic lipid mobilisation. This hypothesis can be tested in the future, for example in flies with disrupted peripheral clocks specifically in the fat body or gut. Independent of whether hemolymph transport lipids originate from *de novo* synthesis in the gut or are mobilised from the fat body, the temporal profile under constant conditions suggests that the clock serves to align transport lipid levels to an optimum time window for anabolism. This hypothesis and possible health impacts need to be tested with tissue-specific genetic manipulations of the clock and underlying metabolic pathways, for which *Drosophila* provides ample tools.

## Author contributions

Kelechi Michael Amatobi: performing metabolite analysis including hemolymph extraction, monitoring food consumption, fly rearing, writing

Ayten Gizem Ozbek-Unal: monitoring locomotor activity, writing

Stefan Schäbler: performing metabolite analysis, analysis troubleshooting

Peter Deppisch: monitoring locomotor activity

Martin J Mueller: designing experiments, consultancy in metabolite analysis

Charlotte Förster: designing experiments, consultancy in chronobiology

Christian Wegener: designing experiments, writing, consultancy in food consumption monitoring, supervision

Agnes Fekete: designing experiments, writing, consultancy in metabolite analysis, supervision

## Acknowledgments

The study was funded by the German Research Foundation (DFG), collaborative research center SFB 1047 “Insect timing”, Project A4 (to AF and MM) in collaboration with projects A1 (to CHF) and B2 (to CW). KMA was supported by a grant of the German Excellence Initiative to the Graduate School of Life Sciences, University of Würzburg; AGOU was funded by a postgraduate scholarship (YLSY) of the Turkish Ministry of National Education. We thank Michel Mayr, Matthias Freund, Markus Krischke, and Maria Lesch for excellent technical help in LC–MS analysis, and Jan von Ackeren, Atinuke Melody Ogunboye and Chidinma Elizabeth Obiekwu for their support in performing the experiments. We are also grateful to Konrad Öchsner for general technical support and Dirk Rieger and Pamela Menegazzi for helpful discussions and advice.

