## Supplemental Figures for "The circadian clock is required for rhythmic lipid transport in the *Drosophila* hemolymph in interaction with diet, photic condition and feeding"

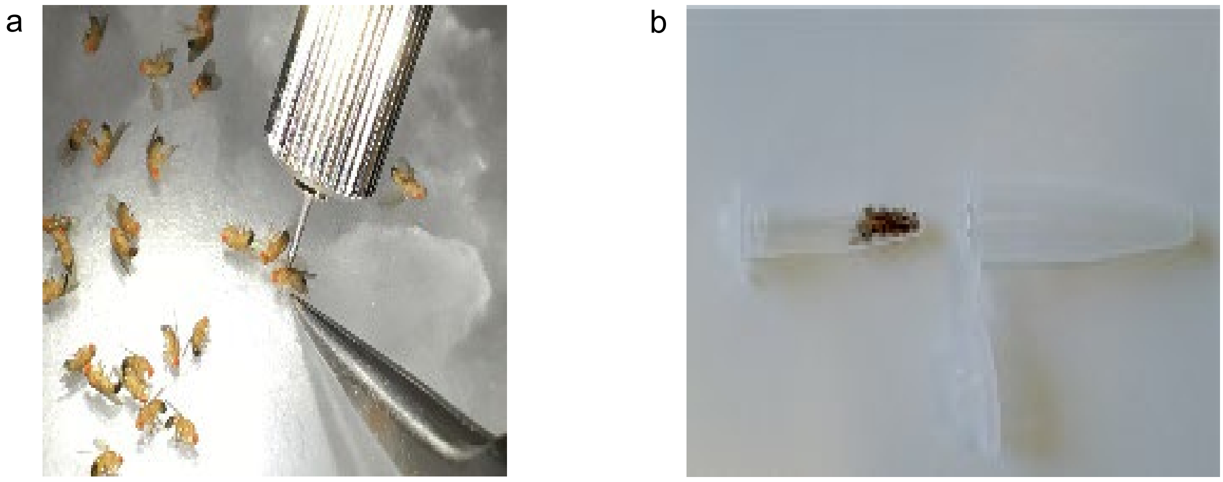


**Suppl Fig S1. Hemolymph sampling.** (a) Six days old male fly were anesthetized on ice and a small incision was made on the metathorax using a tungsten needle. (b) Collection of 20 adult male flies in a 0.5 ml Eppendorf tube with small holes and insertion into 1.5 ml Eppendorf tube.

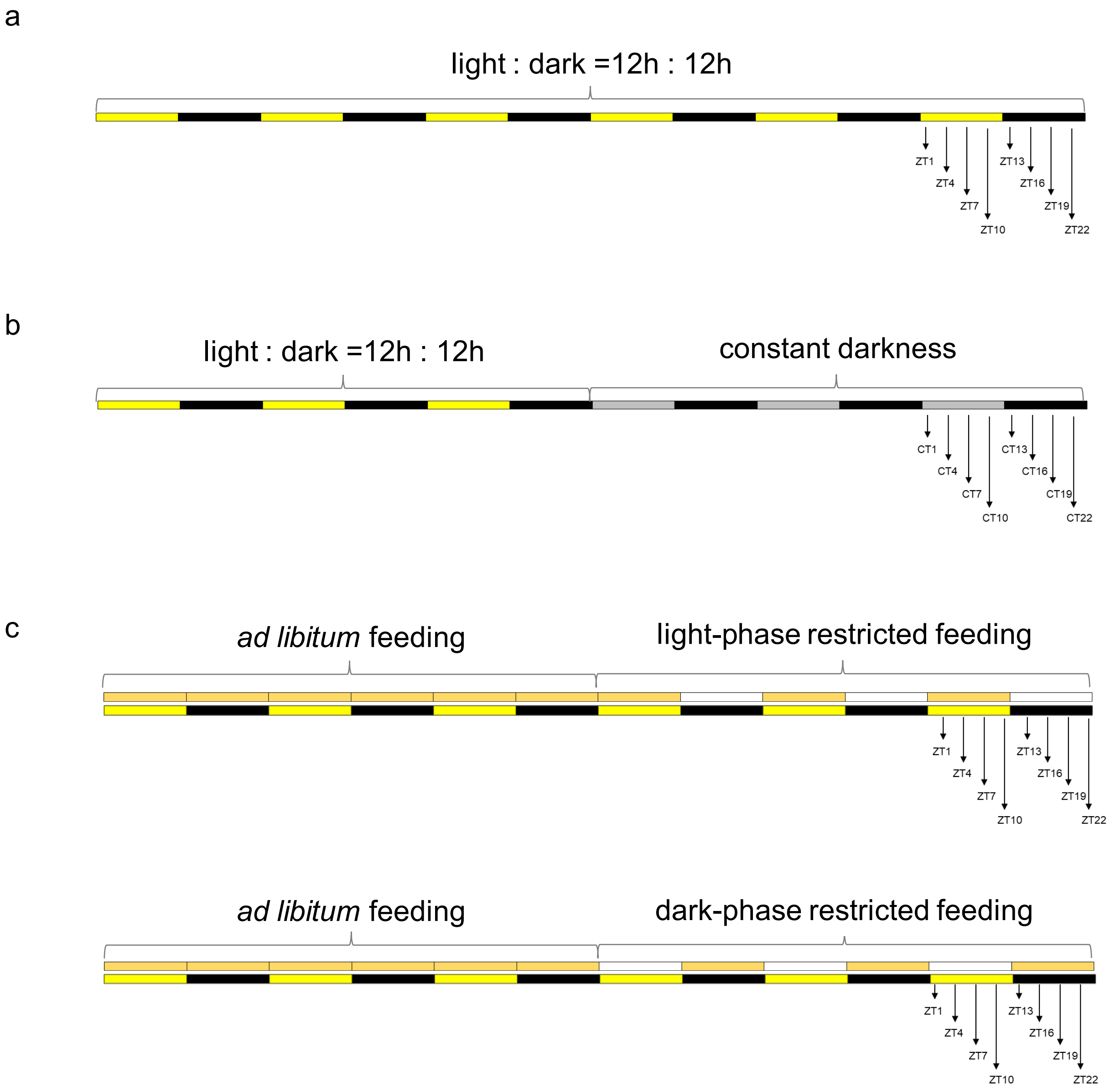


**Suppl Fig S2. Scheme of the experimental treatment and hemolymph extraction.** (a) For studying diel oscillations, male flies were entrained under LD and hemolymph extracted on the 6th day at 8 time points. (b) For monitoring endogen rhythmicity, male flies were entrained for three days under LD cycle before switching to DD and hemolymph extraction on the 6th day at 8 time points. (c) To study the effect of time-restricted feeding, male flies were given food *ad libitum* for three days. Then, food was given only during the light phase (top panel) or only during the dark phase (bottom panel) and sampling of the hemolymph was performed on the sixth day under LD condition. Feeding time is shown in the upper bar, and light conditions in the lower bar. Colour coding: yellow= lights on, black= lights off, orange= 2% agar containing 4% sugar, white = 2% agar.

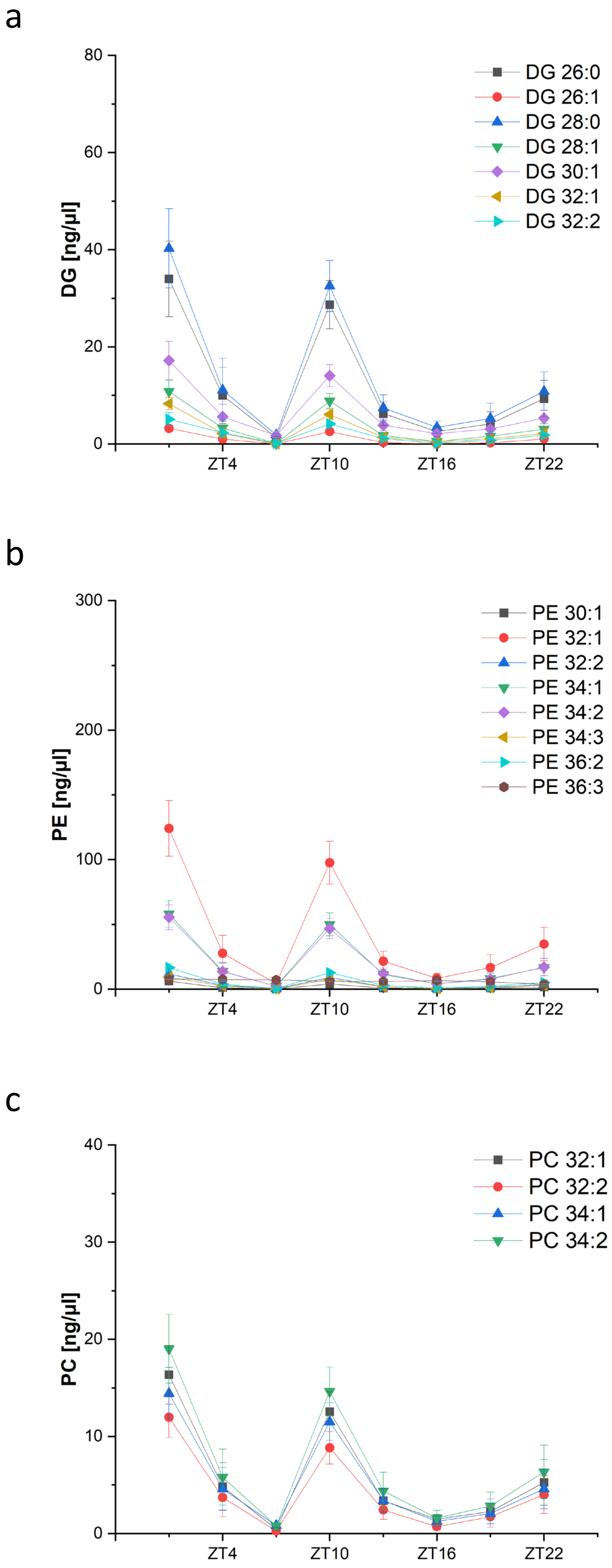


**Suppl Fig S3**. **Temporal pattern of hemolymph lipids in male WT$_{CS}$ flies fed with sugar-only medium is highly similar between individual lipid species.** Levels of detected DG (a), PE (b) and PC species (c) in the hemolmyph of six days old male WT$_{CS}$ flies fed *ad libitum* with sugar-only medium. A lipid species is defined by the lipid class (first two letters), number of acyl carbons (first two numbers) and number of acyl double bounds (last number). Data are presented as mean ± standard error (n=13).

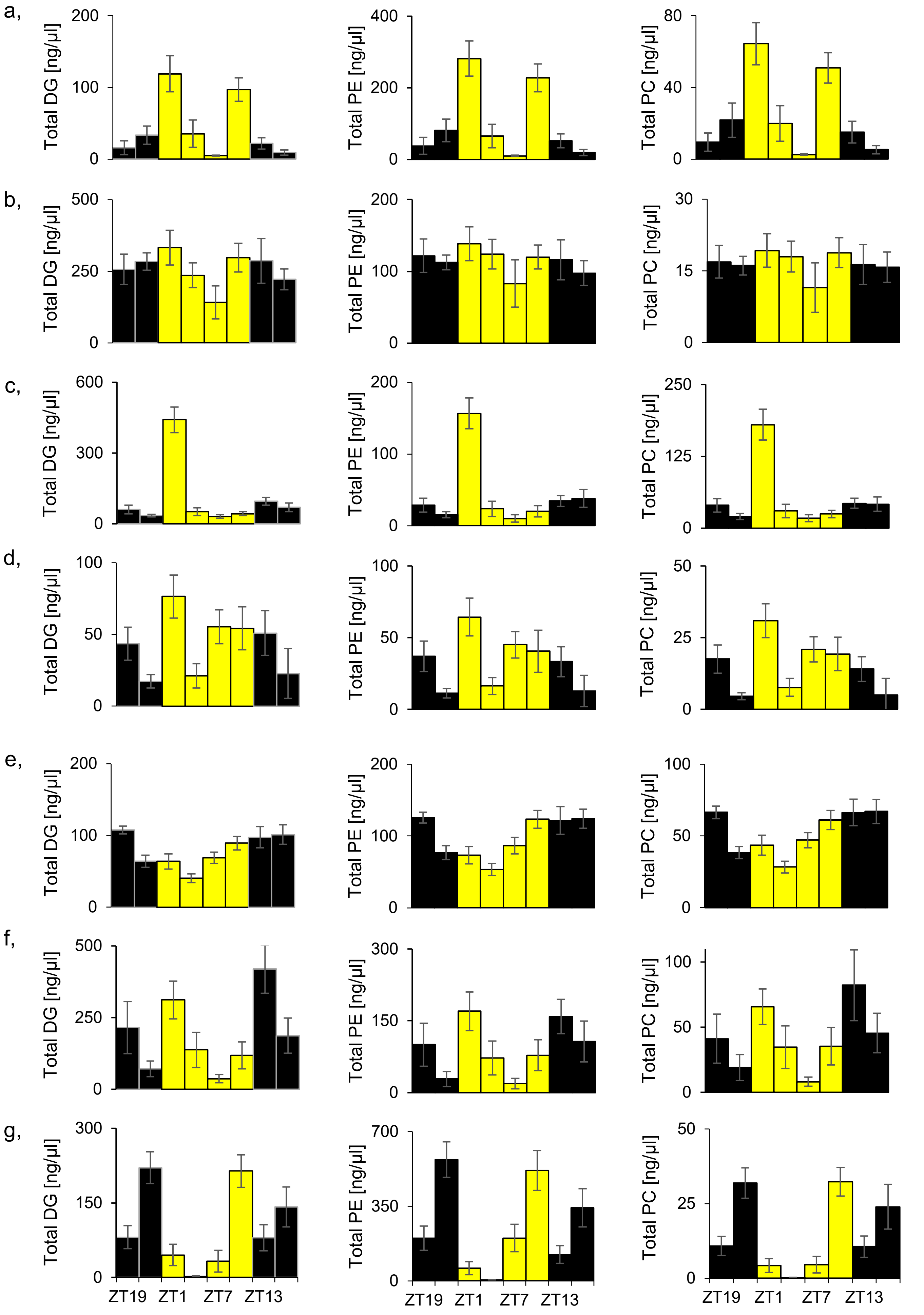


**Suppl Fig S4: Levels of hemolymph lipids in WT$_{CS}$ flies on sugar-only medium depend on time of the day.** Total levels of DGs (left panel), PEs (middle panel) and PCs (right panel) were determined in pooled hemolymph of 20 male WT$_{CS}$ flies using UPLC-TOF-MS. The experiment was repeated seven times independently of each other (a-g). Colour of the bars indicates light conditions (yellow = light-on and black = light off). Data represent mean ± standard error (n=13).

a
b
c

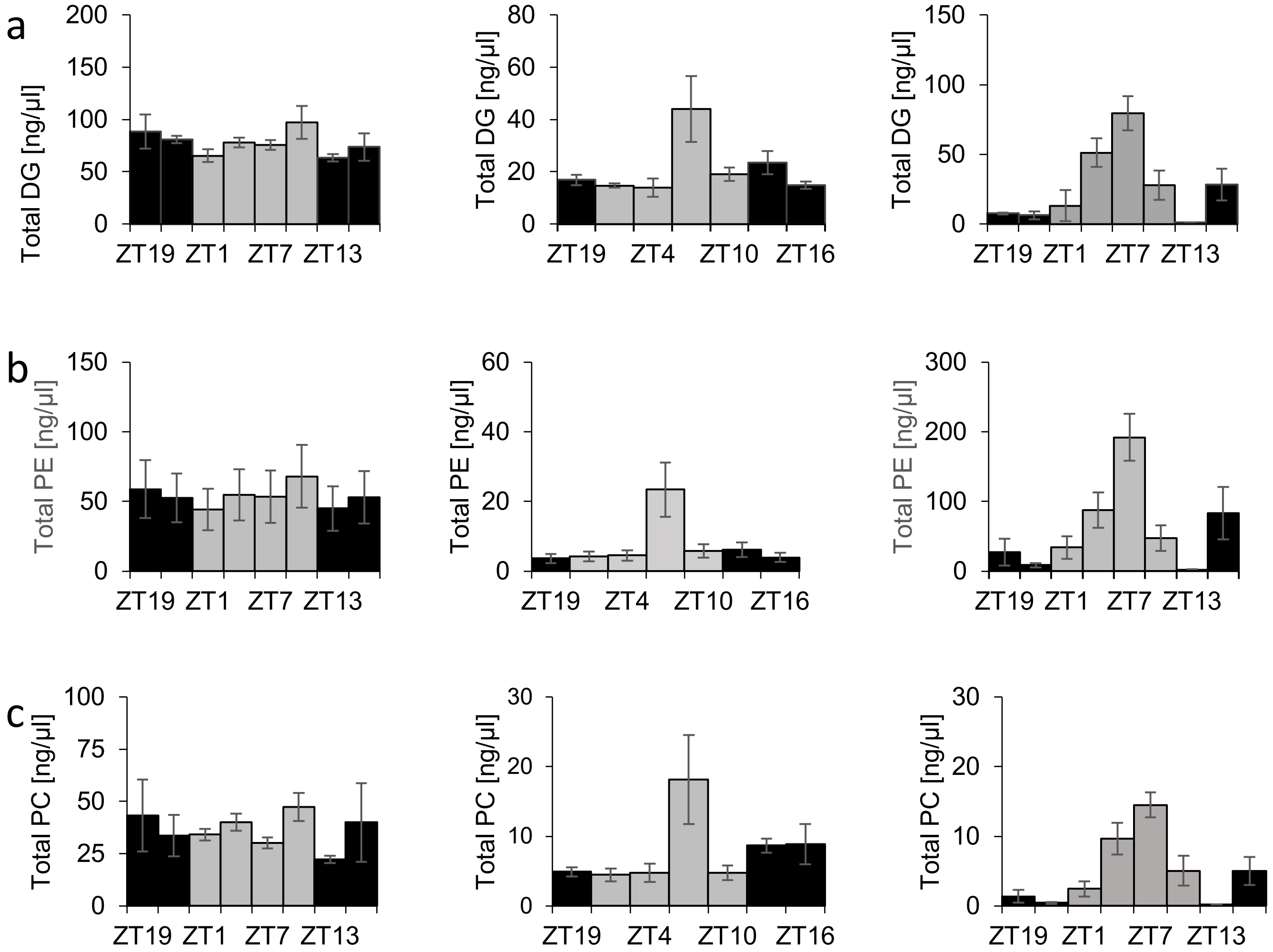


**Suppl Fig S5. Lipid oscillations in the hemolymph of $WT_{CS}$ flies on sugar-only medium weakened and changed under constant darkness.** Three independent experiments showing oscillation profiles of DGs (a), PEs (b) and PCs (c) in the hemolymph of six days old male $WT_{CS}$ fed with sugar-only medium. Flies were synchronized for three days in LD and switched to DD prior to hemolymph sampling every three hours on the sixth day. Light condition is indicated by the colour of the bar (gray= subjective photophase and black = subjective scotophase). Data represent mean ± standard error (n = 13).

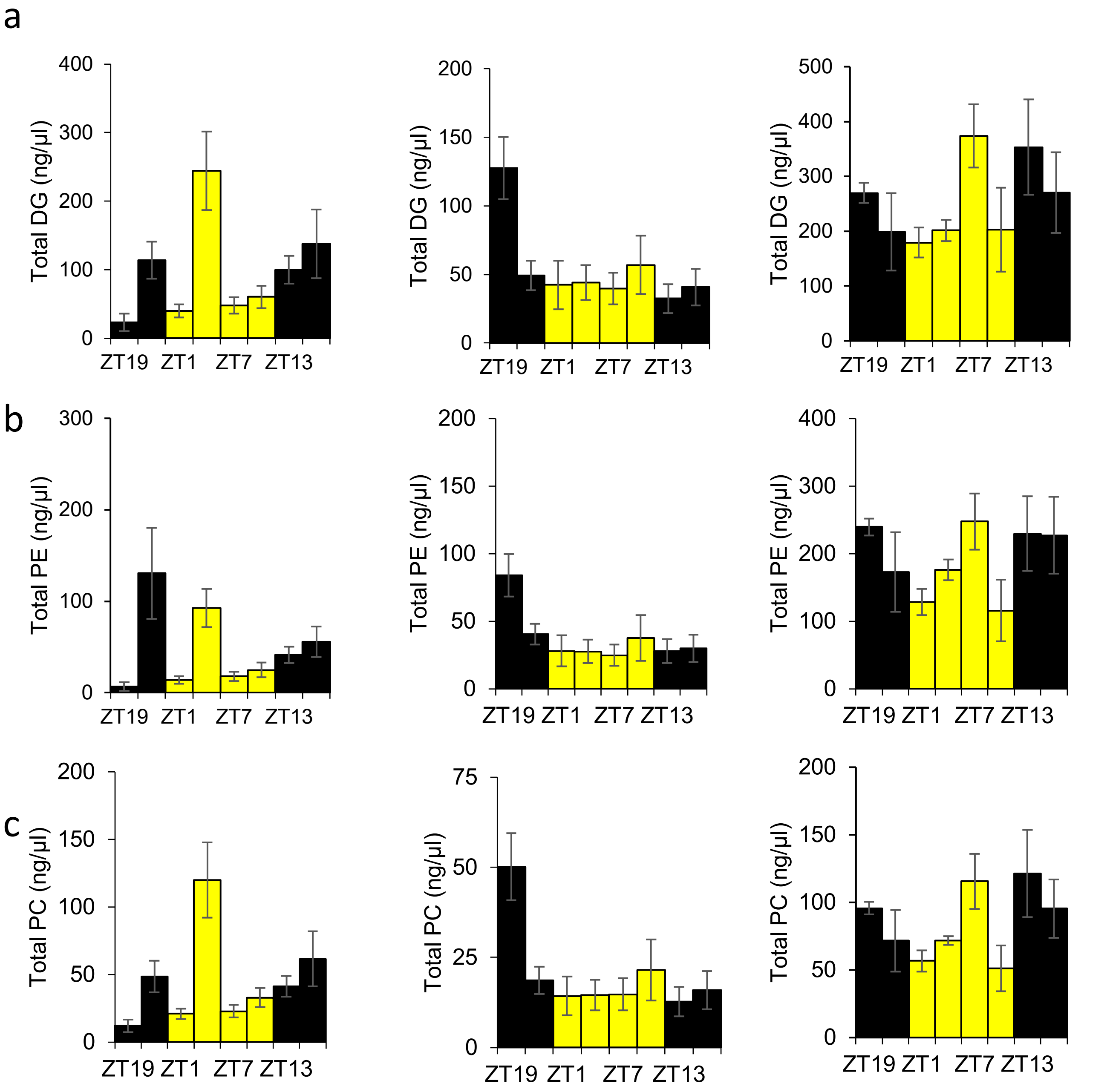


**Suppl Fig S6. Variable oscillations of lipid levels in the hemolymph in *per*$^{01}$ clock mutant flies on sugar-only medium and LD.** Three independent experiments showing oscillation profiles of DGs (a), PEs (b) and PCs (c). Six days old male *per*$^{01}$ flies were fed *ad libitum* with sugar-only medium in LD prior to hemolymph sampling every three hours on the sixth day. Light condition is indicated by the colour of the bar (yellow = lights on and black = lights off). Data are presented as mean ± standard error (n = 13).

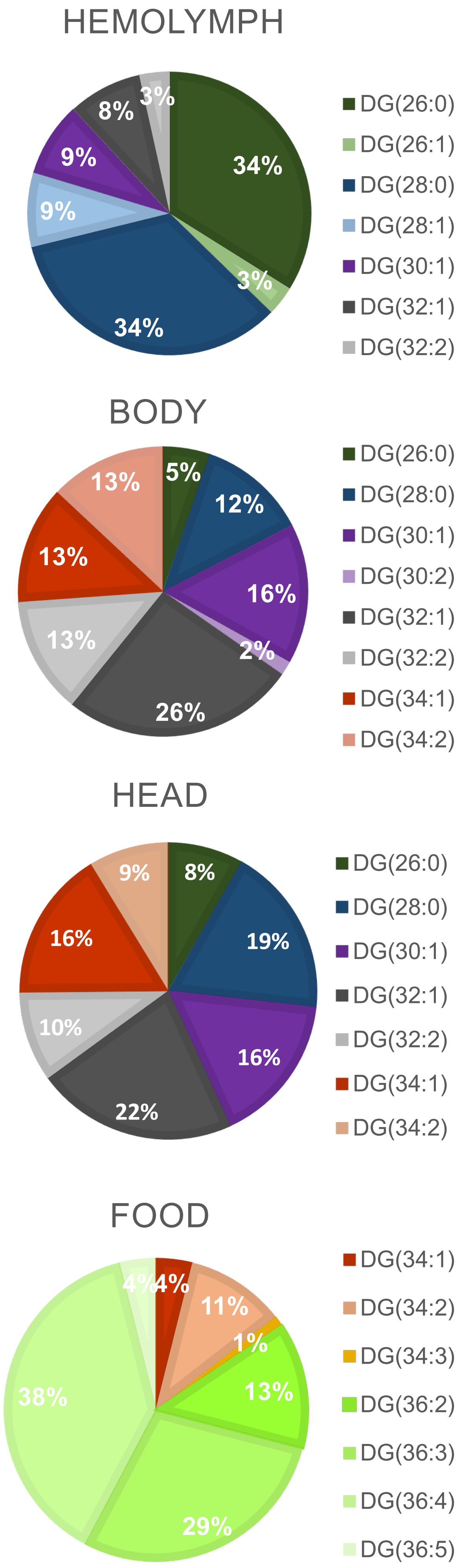


**Suppl Fig S7. Relative species compositions of targeted DGs between hemolymph, body, heads and food in WT$_{CS}$ flies raised on standard medium.** The composition is biased towards lower chain lengths (26-28) in the hemolymph, while body and heads contain mostly longer chain lengths (30-34). Composition of DGs in standard medium differed from the composition in *Drosophila*, since medium-chain DGs were not detectable in standard medium. Lipid species are defined by the lipid class (first two letters), number of carbons (first two numbers) and number of acyl double bounds (last number) of the two fatty acyl chains.

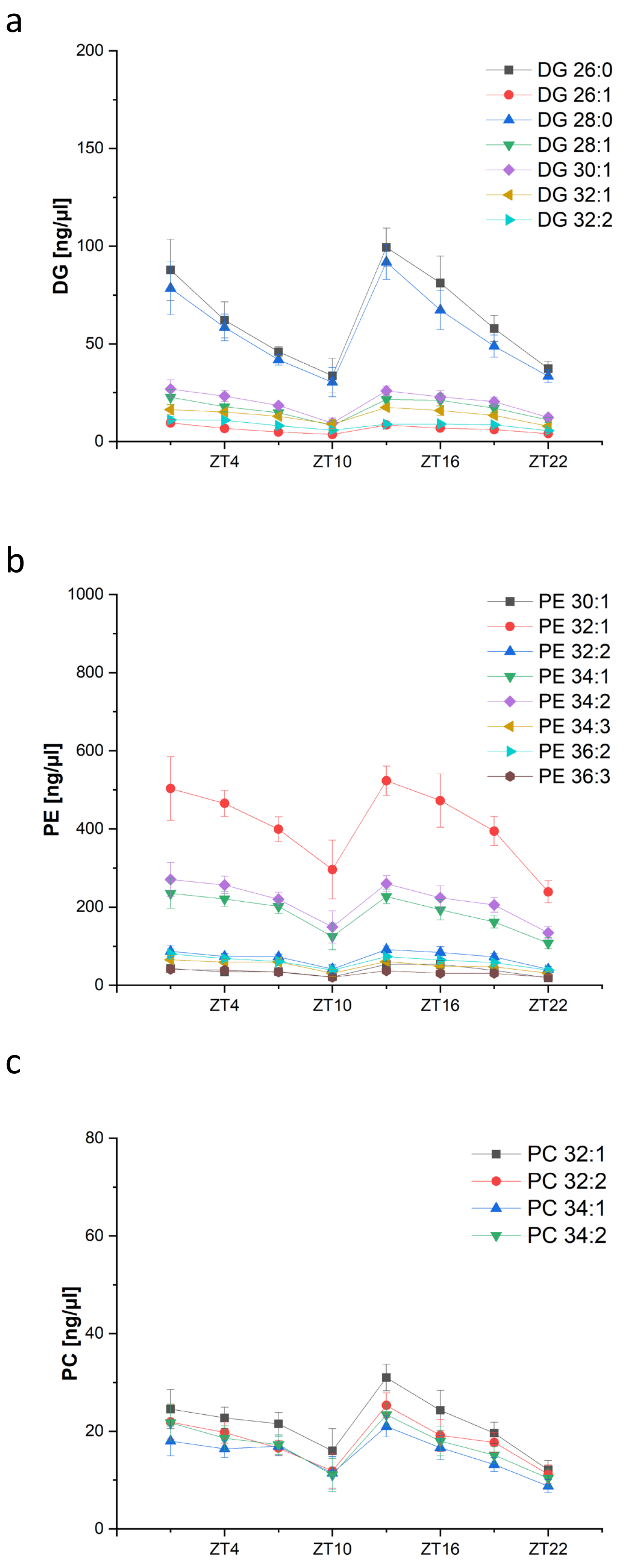


**Suppl Fig S8. Temporal oscillation pattern of hemolymph lipid in presence of dietary lipids remained highly similar between individual lipid species.** Levels of DG (a), PE (b) and PC species (c) in the hemolymph of six days old $WT_{CS}$ male flies fed *ad libitum* with standard medium. Sampling of six days old flies was performed every three hours over a day. A lipid species is defined by the lipid class (first two letters), number of acyl carbons (first two numbers) and number of acyl double bounds (last number). Data are presented as mean ± standard error (n= 13).

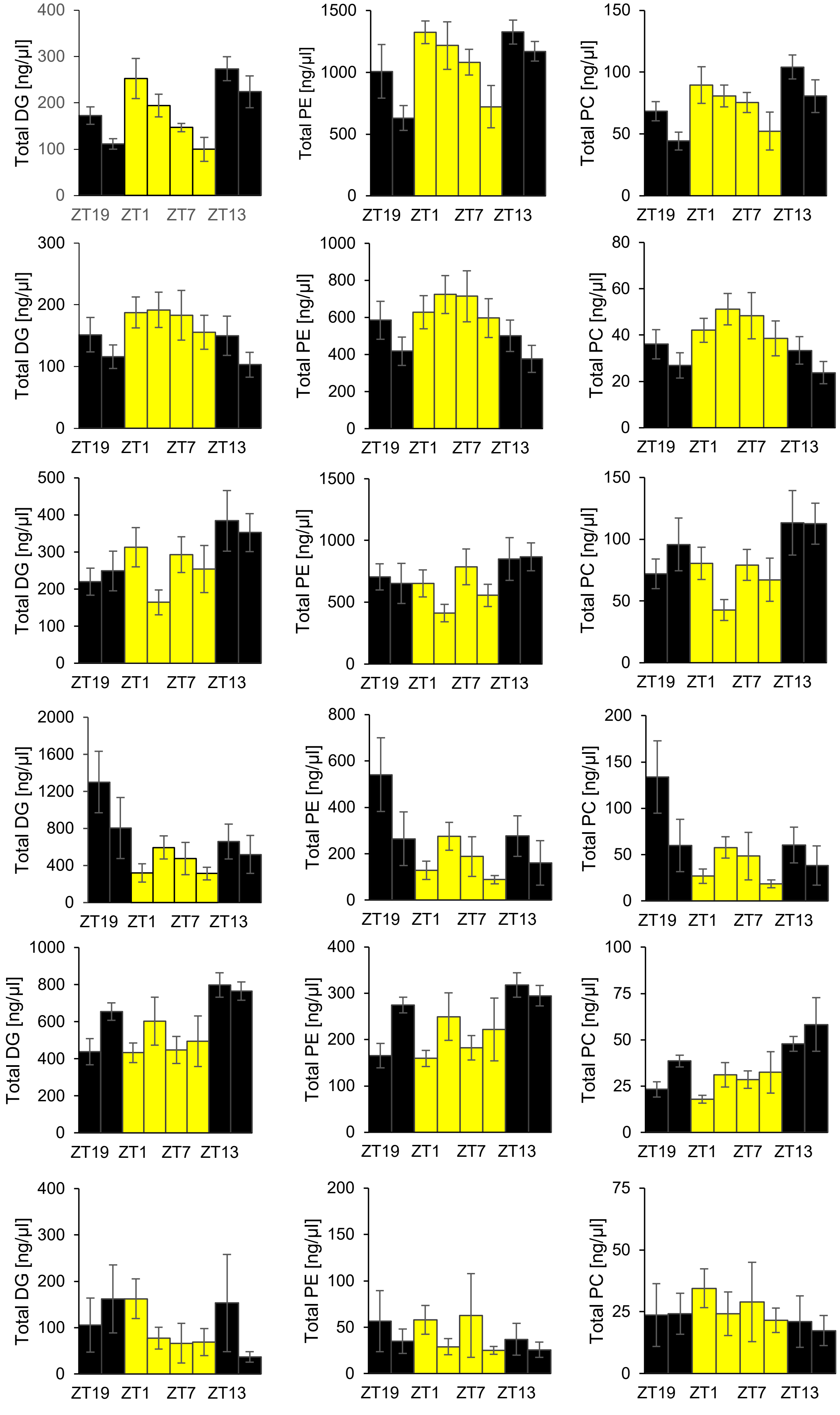


**Suppl Fig S9**. **Dietary lipids mask the daily rhythmicity in circulating lipids.** Total levels of DGs (right panel), PEs (middle panel) and PCs (left panel) in the hemolymph of six days old male $WT_{CS}$ flies fed *ad libitum* with standard medium. Flies were synchronized in LD prior to hemolymph sampling every three hours on the sixth day. Light condition is indicated by the colour of the bar (yellow = lights on, black = lights off). Data are presented as mean ± standard error (n = 13).

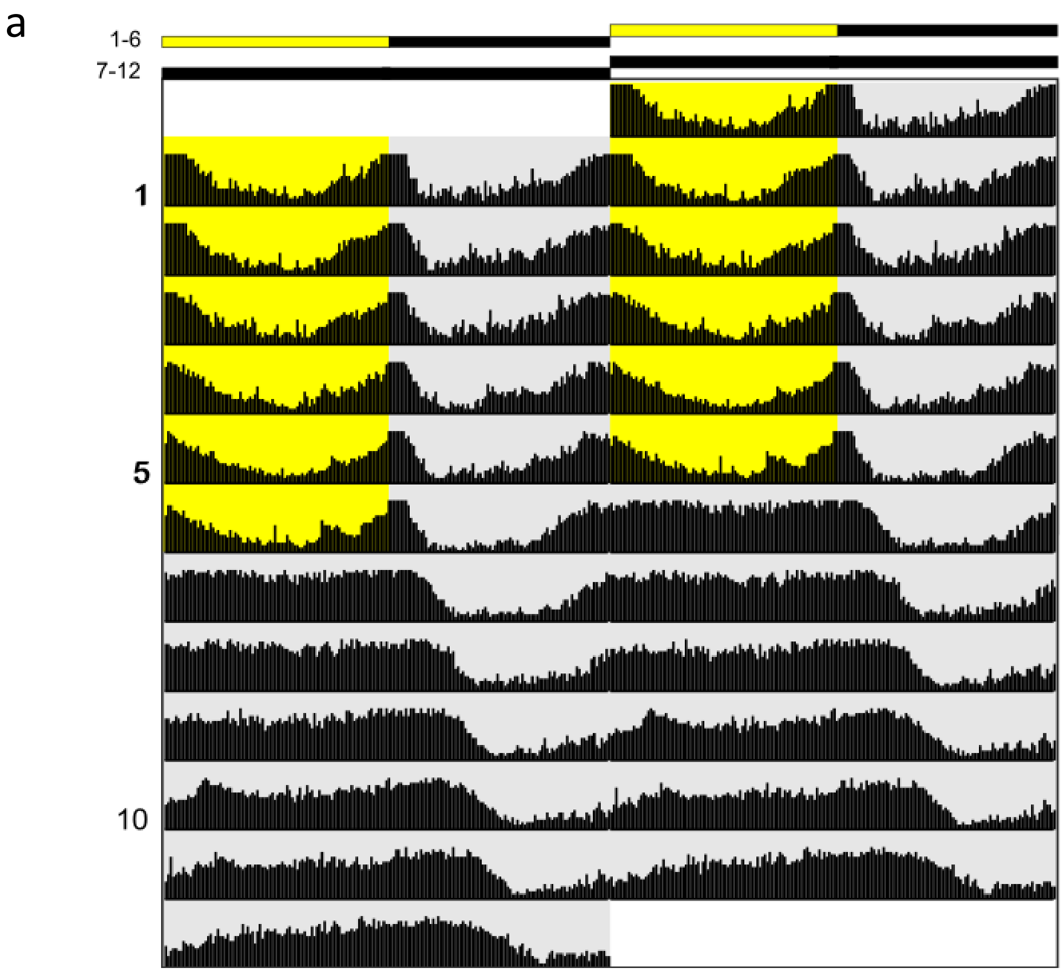


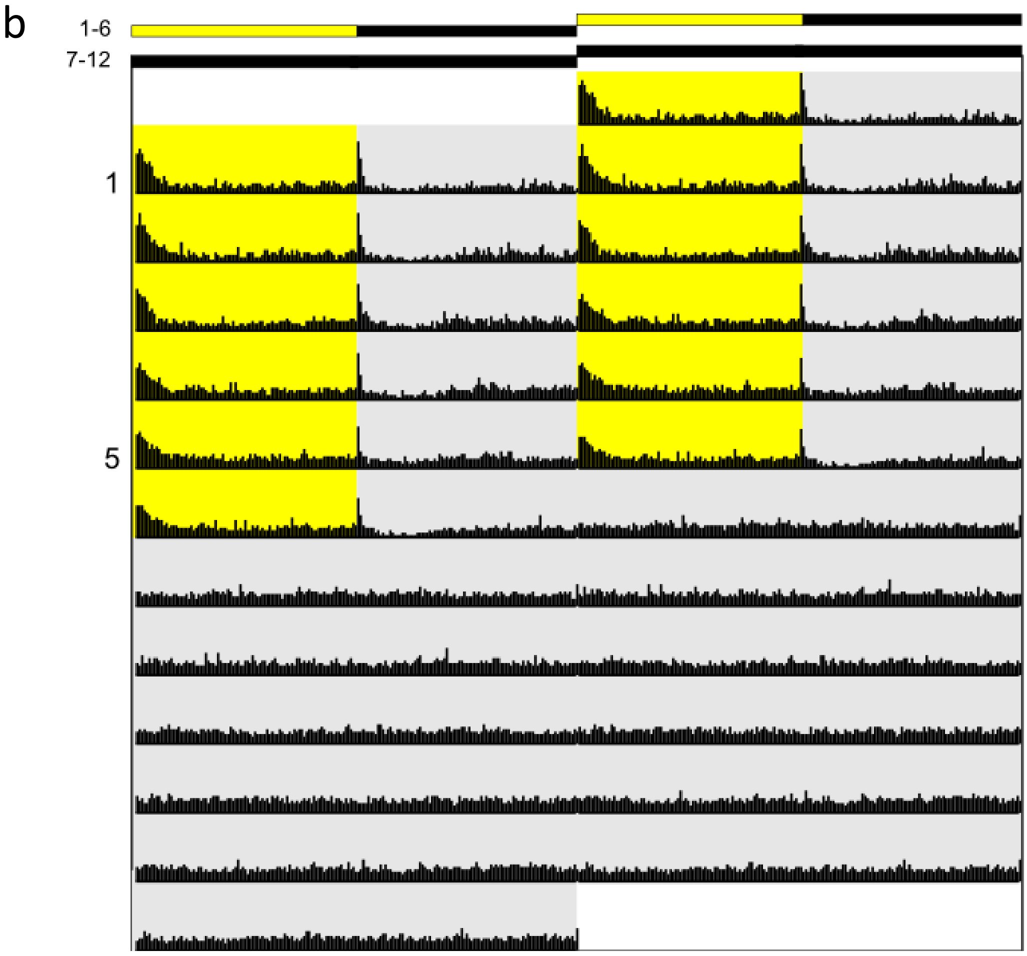


**Suppl Fig S10**. **Average actograms of $WT_{CS}$ (a) and *per*$^{01}$ (b) mutant male flies in LD and DD on standard medium.** Actograms are depicted as double plots. Male flies were recorded for six days in LD and then released to DD for another six days. Light phases are highlighted by a yellow background, dark phases are highlighted by a gray background. $WT_{CS}$: N=29, *per*$^{01}$: n= 32.

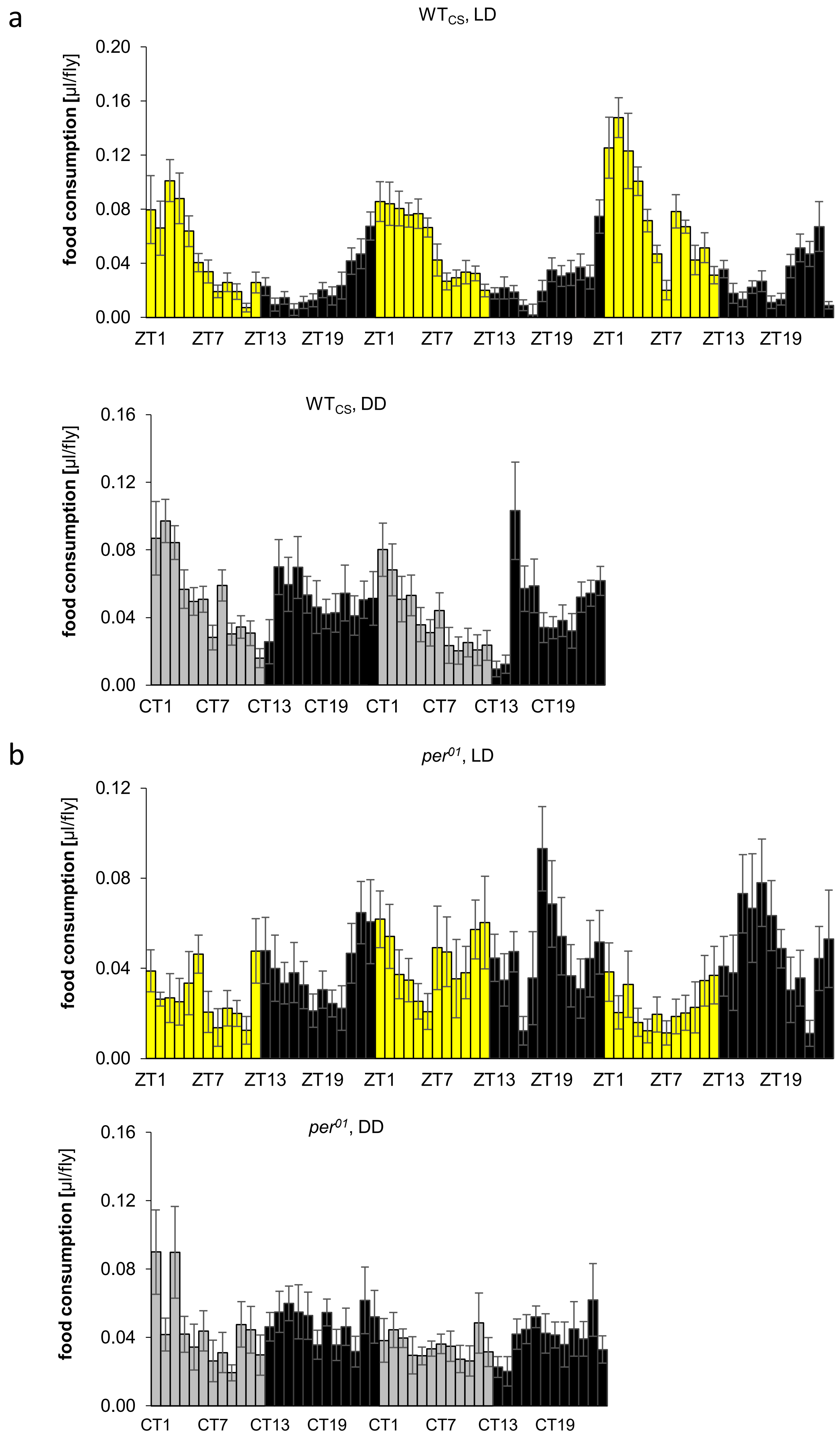


**Suppl Fig S11. Diel rhythmicity of food consumption is dampened in *per$^{01}$* clock mutants on standard medium.** Feeding of male WT$_{CS}$ flies under LD (top panel in a) and DD (bottom panel in a) and of *per$^{01}$* flies under LD (top panel in b) and DD (bottom panel in b) were investigated from fourth to sixth day under LD and from first to second day in DD after three days entrainment in LD. Food consumption was monitored by a modified CAFE assay. The food capillary was filled with 3.6% (w/w) yeast, 5% (w/w) sucrose containing 0.3% (v/v) FD&C Blue No 1 (E133) blue. Food was changed every two days between ZT20 and ZT22. Light condition is indicated by the colour of the bars (LD: yellow = lights on, black = lights off; DD: light gray = subjective photophase, dark gray = subjective scotophase). Data are presented as average of two independent experiments ± standard error (N=2, n=10).

a

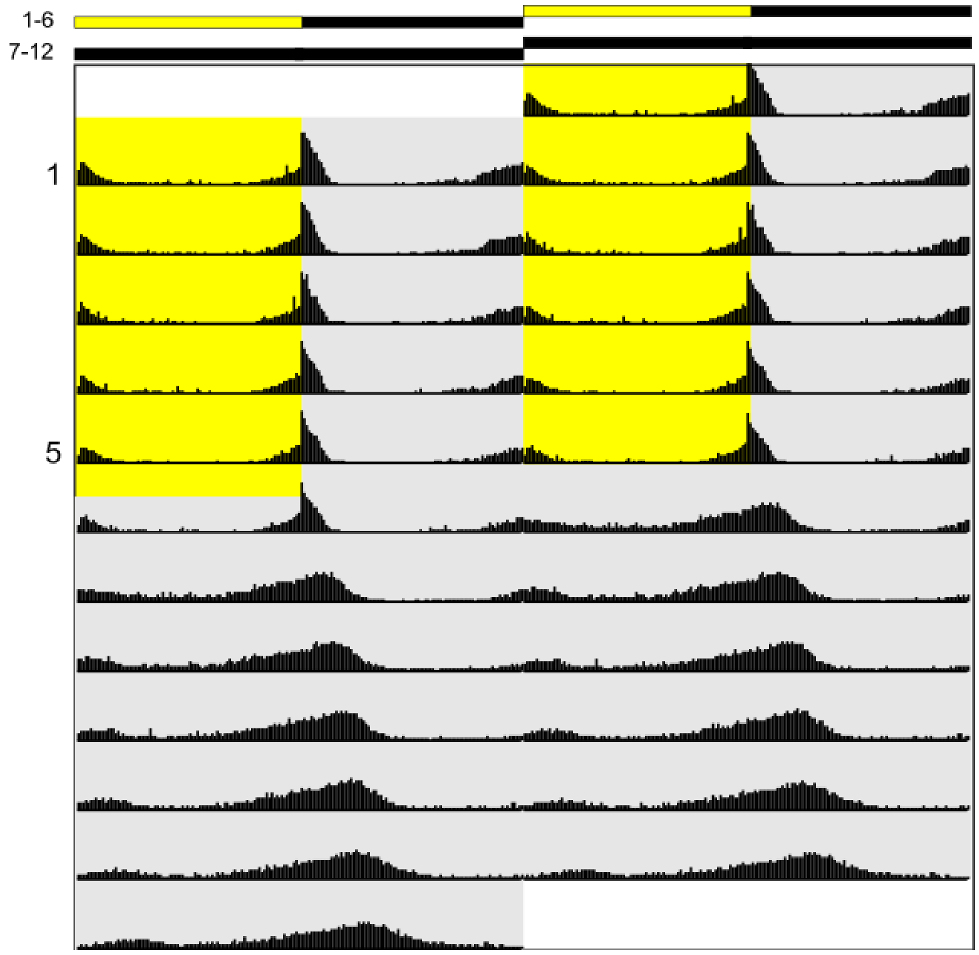


b

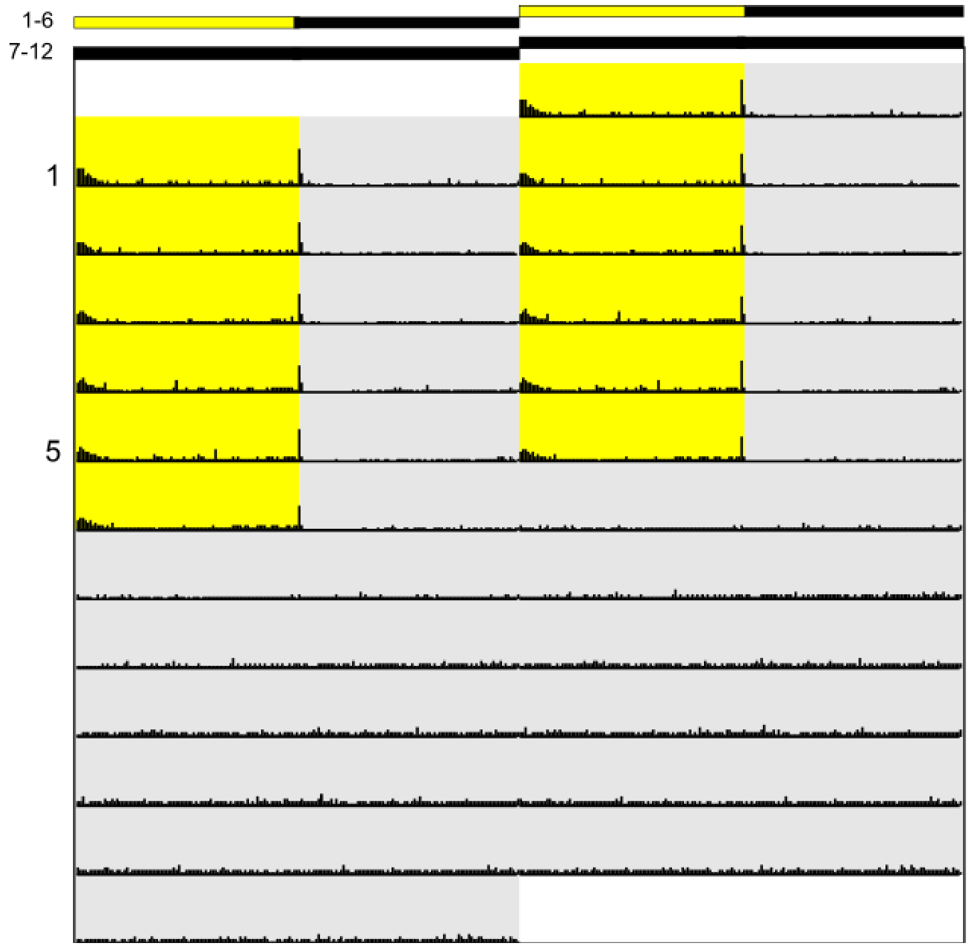


**Suppl Fig S12 Average actograms of a) WT$_{CS}$ and b) *per*$^{01}$ mutant male flies in LD and DD on sugar-only medium** Actograms are depicted as double plots. Male flies were recorded for six days in LD and then released to DD for another six days. Flies were fed *ad libitum* with 5% (w/w) sucrose and 3% agar. Light phases are highlighted by a yellow background, dark phases are highlighted by a gray background. WT$_{CS}$: n= 32; *per*$^{01}$: n= 32.

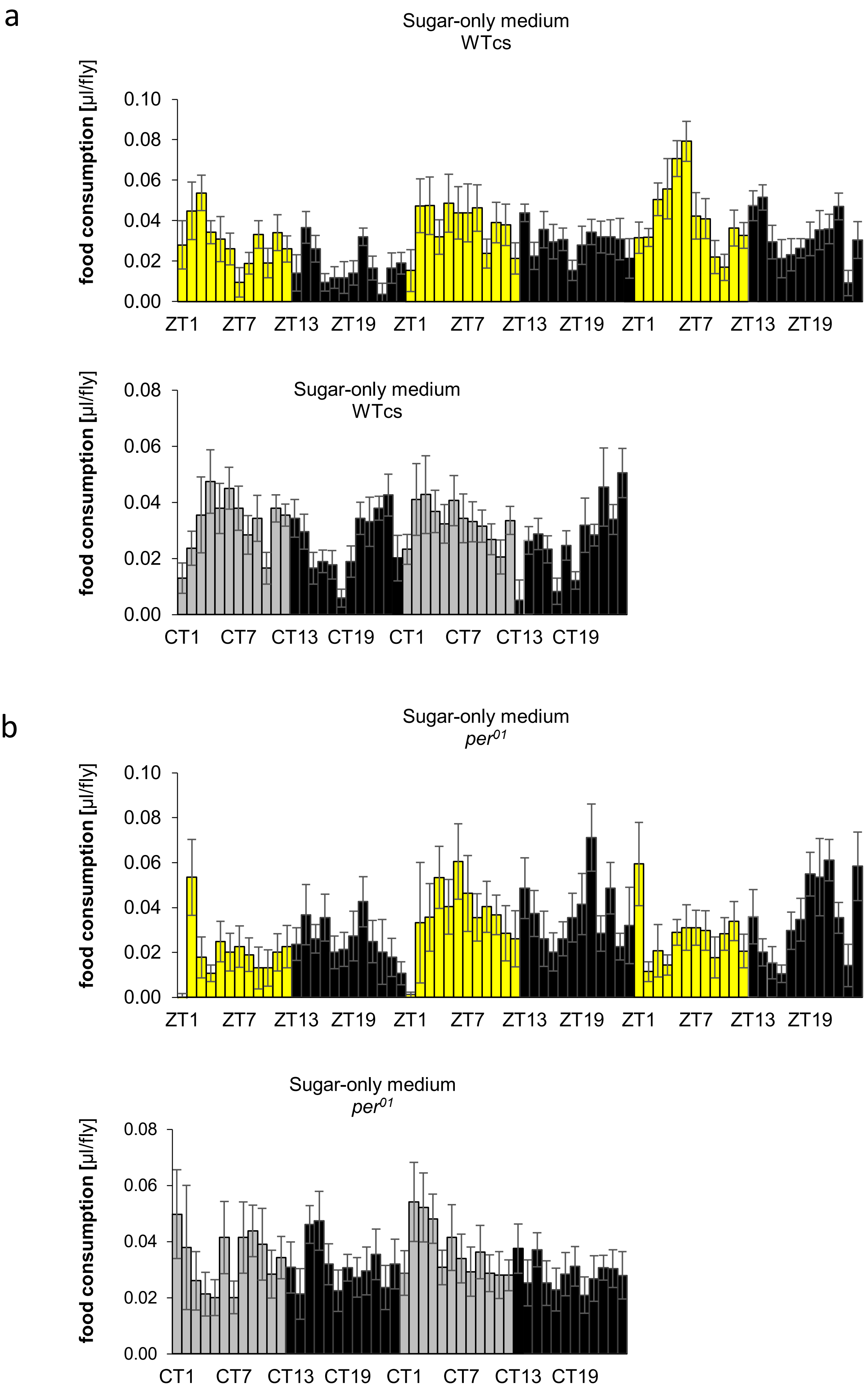


**Suppl Fig S13. Diel rhythmicity of food consumption is dampened in $WT_{CS}$ wild type flies on sugar-only medium.** Food consumption of $WT_{CS}$ wild type flies under LD (top panel in a) and DD (bottom panel in a), and of *per*$^{01}$ flies under LD (top panel in b) and DD (bottom panel in b) was investigated from fourth to sixth day under LD and from first to second day in DD after three days entrainment under LD. Food consumption was monitored with a modified CAFE assay. Food capillary was filled with 5% (w/w) sucrose containing 0.3% (v/v) FD&C Blue No 1 (E133) blue. Food was changed every two days between ZT20 and ZT22. Light conditions are indicated by the colour of the bars (LD: yellow = lights on, black = lights off; DD: light gray = subjective photphase, dark gray = subjective scotophase). Data are presented as average of two independent experiments ± standard error (N=2, n=10).
